# Exploiting functional regions in the viral RNA genome as druggable entities

**DOI:** 10.1101/2024.08.11.607475

**Authors:** Dehua Luo, Yingge Zheng, Zhiyuan Huang, Zi Wen, Lijun Guo, Yingxiang Deng, Qingling Li, Yuqing Bai, Shozeb Haider, Dengguo Wei

## Abstract

RNA-targeting compounds or small interfering RNAs (siRNAs) offer a potent means for controlling viral infections. An essential prerequisite to their design depends on identifying conserved and functional viral RNA structures in cells. Techniques that probe RNA structures *in situ* have been developed recently including SHAPE-MaP, which has been helpful in analyzing the secondary structures of RNA. In this study, we report on the application of SHAPE-MaP to the Porcine Epidemic Diarrhoea Virus (PEDV) RNA genome to categorize different functional regions including potential quadruplex forming sequence and target sites of siRNA. Our results show that these structures can be exploited to inhibit viral proliferation and that SHAPE-MaP is an effective method to the identification of secondary structures in RNA genomes.

## INTRODUCTION

RNA-targeting drugs are expected to address the challenges posed by "undruggable" protein targets and significantly expand the landscape of targetable macromolecules(Disney, 2019; Kovachka et al., 2024; Li and Wang, 2019; Ursu et al., 2020; Warner et al., 2018). The U.S. Food and Drug Administration (FDA) has approved six siRNAs(Tang and Khvorova, 2024), nine antisense oligonucleotides (ASOs)(Bennett, 2019; Chen et al., 2024; Crooke et al., 2021), and the RNA-targeting compound Risdiplam(Ratni et al., 2021; Sheridan, 2021) for treating complicated diseases such as spinal muscular atrophies. The selection of RNAs that are functional, conserved and accessible to be targeted *in vivo* is fundamental to RNA-targeting drug development. Unlike RNAs *in vitro*, the folding of RNAs in cells is affected by the intricate intracellular environment(Manfredonia et al., 2020; Zhang et al., 2021). Thus, *in situ* probing for the accessibility of viral RNAs by antivirals is essential for the identification of targetable RNAs.

In recent years, the integration of chemical probing methods and high-throughput sequencing technologies has been employed to analyse the structure of the viral RNA genome in infected cells(Boerneke et al., 2019; Cao et al., 2021; Flynn et al., 2016; Huber et al., 2019; Huston et al., 2021; Lan et al., 2022; Li et al., 2018; Manfredonia et al., 2020; Mauger et al., 2015; Pollom et al., 2013; Siegfried et al., 2014; Smola et al., 2016; Yang et al., 2021; Zhang et al., 2021; Ziv et al., 2020). These methods have confirmed the widespread presence of structurally organized RNA elements within the viral genome, which are composed of various types of RNA structures such as stem-loop structures (with double-helical stems), RNA single-strand, RNA pseudoknot and RNA G-quadruplex (G4), collectively forming a complex and dynamic global genome architecture(Boerneke et al., 2019; Kwok et al., 2016). Of these, RNA G4 has shown considerable promise because of its high stability and ability for modulation by small molecules(Fang et al., 2023; Fleming et al., 2016; Kwok et al., 2016; Ruggiero and Richter, 2018; Wang et al., 2016; Zhao et al., 2021). Furthermore, RNA interference (RNAi) has been demonstrated to inhibit expression of genes and viral replication(Bennett, 2019; Bowden-Reid et al., 2023; Piasecka et al., 2020; Qureshi et al., 2018; Sagan et al., 2010; Tang and Khvorova, 2024; Westerhout and Berkhout, 2007). In spite of the advances, identifying RNA elements or regions that serve as targets within complex genomic structural contexts remains challenging.

Selective 2’-hydroxyl acylation and primer extension mutational profile (SHAPE-MaP) stands out in the *ex vivo* RNA structure probing technologies for its ability to quantify nucleotide flexibility and solvent accessibility at single-nucleotide resolution(Lorenz et al., 2016; Siegfried et al., 2014; Smola et al., 2016),(Lucks et al., 2011). In short, SHAPE reagents like NAI selectively modify flexible, unpaired 2’-OH groups in RNA, and these modifications are detected as mutations during reverse transcription, enabling precise mapping of RNA secondary structures through sequencing (Siegfried et al., 2014). In the viral long-stranded RNA genome, SHAPE reactivities are used as constraints in the prediction of RNA structures based on nearest-neighbour rules(Deigan et al., 2009; Mathews et al., 2004), which may result in thousands of possible conformations. The pairing probability of each nucleotide derived from SHAPE reactivities was subsequently used to calculate Shannon entropy. Regions with high Shannon entropy may adopt alternative conformations, while those with low Shannon entropy correspond to either well-defined RNA structures or persistently single-stranded regions (MATHEWS, 2004; Siegfried et al., 2014). Therefore, SHAPE reactivity provides information about the pairing probability and accessibility of nucleotides, while Shannon entropy reflects the likelihood of alternative conformations in a given region. RNA motifs with low SHAPE reactivity and low Shannon entropy are assumed to be well fold structures and potential targets(Boerneke et al., 2019). In this study, we categorized RNAs with distinct features in the viral genome according to the SHAPE reactivity and Shannon entropy, and investigate their potential as targets (Figure 1).

**Figure 1.**
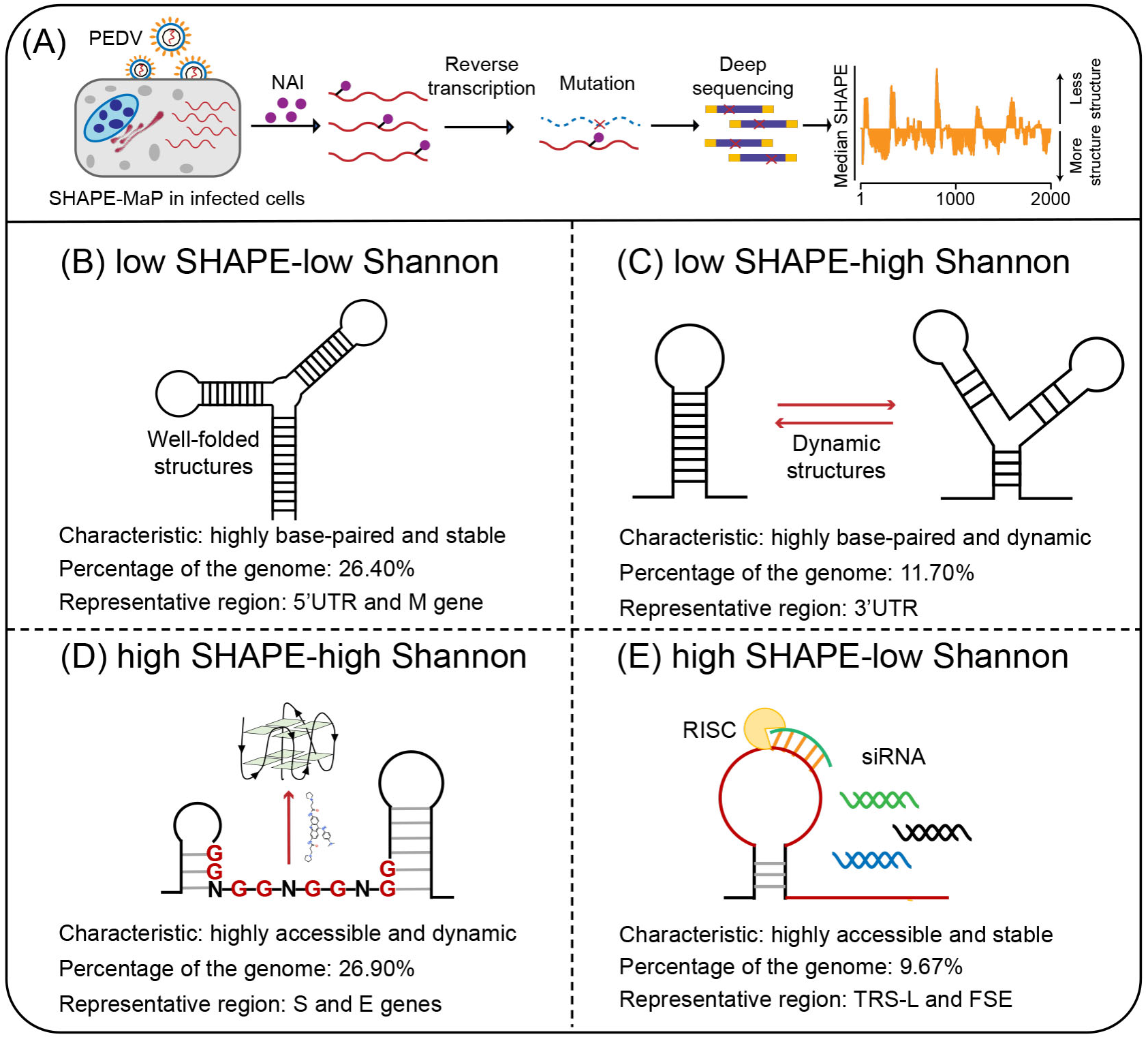
Four regions with different characteristics in the PEDV genome. **(A)** Schematic of SHAPE-MaP for probing the RNA structure of the PEDV genome *in situ*. **(B)** Well-folded regions (low SHAPE reactivity and low Shannon entropy; 26.40% of genome). These regions represent stably folded RNA structures with minimal conformational flexibility, likely serving as structural scaffolds or functional elements in viral replication. **(C)** Dynamic structured regions (low SHAPE reactivity and high Shannon entropy; 11.70% of genome). These conformationally plastic domains likely mediate regulatory switches between alternative secondary structures during infection. **(D)** Dynamic unpaired regions (high SHAPE reactivity and high Shannon entropy; 26.90% of genome). These regions are prone to form non-canonical nucleic acid structures (e.g., G-quadruplexes), which can be stabilized by small-molecule ligands to inhibit viral replication. **(E)** Persistent unpaired regions (high SHAPE reactivity and low Shannon entropy; 9.67% of genome). These regions are more accessible for siRNA binding, facilitating recruitment of Argonaute proteins and Dicer to form the RNA-induced silencing complex (RISC) for targeted cleavage.

Porcine epidemic diarrhoea virus (PEDV) is an α-coronavirus with a positive single-stranded RNA genome of 28 kb(Jung et al., 2020; Lee, 2015). It results in severe diarrhoea and vomiting, leading to the death of ∼90 percent of infected piglets. The spread of PEDV is world-wide and its infection is of primary concern in pig husbandry(Lee, 2015). The epizootic re-emergence of PEDV has also been reported to infect human intestinal cells and displays potential susceptibility to cross-species infection(Niu et al., 2023).

In this study, we adapted the SHAPE-MaP method to detect the *in situ* structure of the PEDV RNA genome in infected cells. More specifically, the combination of SHAPE reactivity and Shannon entropy categorized the PEDV RNA genome into different functional types (Figure 1), based on the distinctive profiles, including the 5’ untranslated region (5’ UTR), frameshift stimulatory element (FSE), and 3’ untranslated region (3’ UTR). We particularly focused on guanine-rich regions, which could potentially fold into a stable G4 structure and sites suitable for binding small interfering RNA (siRNA). Our results show that the folding of G4 in a dynamic single-stranded region with high SHAPE reactivity and high Shannon entropy inhibited viral proliferation, while siRNA targeting persistent single-stranded regions with high SHAPE reactivity and low Shannon entropy exhibited greater success rates in inhibiting viral replication. Finally, this work proposes a strategy for selecting targetable RNAs based on the *in situ* folding characteristics of the viral genome.

## RESULTS

### Mapping RNA structures in the PEDV genome

SHAPE-MaP was used to characterize the structural features of the PEDV RNA genome in infected cells (Figure 1A). The high correlations of mutational signatures for biological replicates (*ex vivo* SHAPE R^2^=0.955; Figure 1—figure supplement 1A) and the high mutation rate with NAI treatment (Figure 1—figure supplement 1B) indicate the high quality of the sequencing data and the experimental workflow. Hence, two replicates were combined for downstream analyses.

As the 5’UTR with conserved functions across coronaviruses has been extensively reported(Chen and Olsthoorn, 2010; Madhugiri et al., 2018; Yang and Leibowitz, 2015), we first examined the structure of this region in the PEDV genome for its stability *ex vivo*. The PEDV 5’ UTR structure modelled with SHAPE reactivity constraints contains four stem loops (SL1, SL2, SL4, and SL5; Figure 1—figure supplement 1C) similar to other alpha-coronaviruses(Chen and Olsthoorn, 2010; Madhugiri et al., 2018; Yang and Leibowitz, 2015). Our SHAPE dataset showed that ∼95.6% of highly reactive bases (SHAPE reactivity ≥ 0.7) are not involved in canonical base pairing or are located adjacent to helical termini or bulges/loops (Figure 1—figure supplement 1C). This result suggested that our dataset is reliable as a constraint for predicting the intracellular genomic structure of PEDV.

To characterize the intracellular folding features of the PEDV genome, median SHAPE reactivities and Shannon entropies were calculated in sliding, centered windows (window = 50 nt, step = 1 nt), and a full-genome SHAPE structural map was constructed accordingly (Figure 2). The global median SHAPE reactivity of the genome is 0.233. Specifically, within the 1- 8000 nt region at the 5’ end, the median reactivity is 0.173; it increases to 0.268 in the middle segment of the genome (8000-24000 nt) and then decreases to 0.194 at the 3’ end (24000-28000 nt). These results indicate that the folding of PEDV genomic RNA within infected cells is nonuniform, with the 5’ and 3’ ends being more compactly structured, while the central region is more flexible (Figure 2).

**Figure 2.**
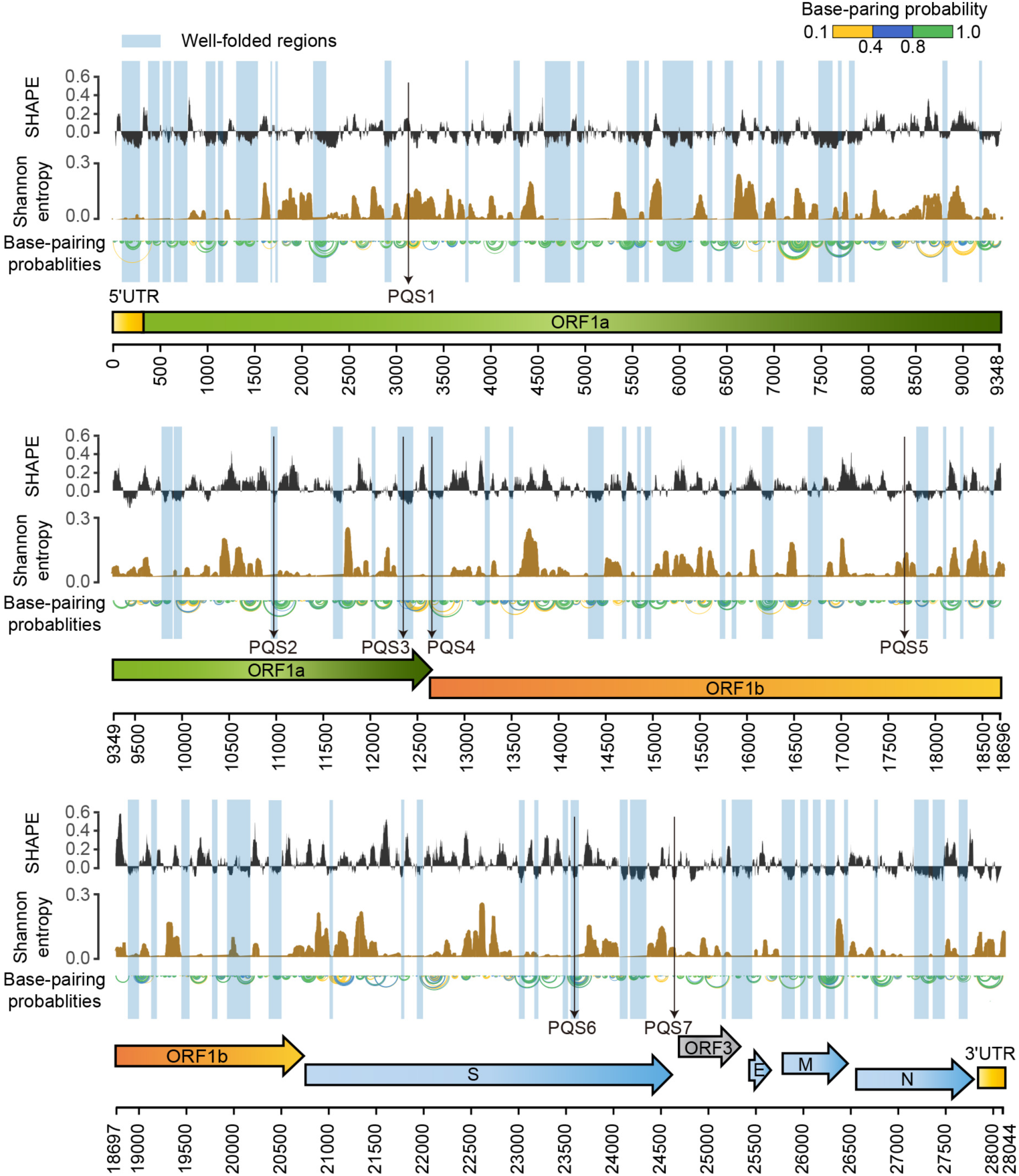
SHAPE structural map of the PEDV genome in infected cells. From top to bottom: SHAPE reactivity, Shannon entropy, base-pairing probabilities, translation reading frames are indicated by arrows, and genomic coordinates are marked at the bottom. Well-folded regions with low SHAPE reactivity and low Shannon entropy are shaded in blue. Potential G4 forming sequences (PQSs) are marked with a black arrow.

Within the coding regions of the ORF1a, ORF3, M, and N genes, both SHAPE reactivity and Shannon entropy are below the global median, suggesting that these regions are stably folded in the genome (Figure 2—figure supplement 1). In contrast, the coding regions of the S and E genes exhibit more relaxed and variable conformations, with both SHAPE reactivity and Shannon entropy exceeding the global median (Figure 2—figure supplement 1). Among the known functional regions, the 5’ UTR (position: 1-310 nt), which is involved in translation initiation and genome packaging, exhibited lower SHAPE reactivity and lower Shannon entropy compared to the global median, indicating a well-folded overall structure. The frameshifting stimulatory element (FSE) (position 12605-12690), which is involved in mediating ribosomal frame shifting, exhibited high SHAPE reactivity and low Shannon entropy values. The 3’ UTR (position 27702-28044 nt), which is responsible for genome cyclization, mRNA stability, and initiation of negative-strand synthesis, showed low SHAPE reactivity and high Shannon entropy values. These functional regions in the PEDV genome exhibiting distinct SHAPE reactivity and Shannon entropy profiles suggest that these regions may adopt different structural folding patterns to perform specific functions (Figure 1).

### Functional categorization of PEDV genome reveals potentially targetable RNA

To further analyse other functional regions through SHAPE reactivity and Shannon entropy, we combined these two parameters for the classification of structural features (see Methods). Low SHAPE regions were defined as windows in which >75% of the bases exhibited SHAPE reactivities below the global median (calculated on the full PEDV genome), and low Shannon entropy regions were defined according to the same rule, following Manfredonia’s method (Manfredonia et al., 2020). Windows in which >75% of their bases exhibiting values above the global median were picked as high SHAPE or high Shannon regions. Finally, four categories of RNAs in the PEDV genome were defined according to the relative values of SHAPE reactivity and Shannon entropy. (Figure 1, Table S2 and Table S3).

Regions with high base-pairing rates (low SHAPE) accounted for 38.10% of the PEDV genome, including the 5’ UTR and 3’ UTR (Figure 1B-C). Of these the low SHAPE-low Shannon regions (26.40%) maintain stable conformations throughout the viral life cycle and are assumed to represent the well-folded skeleton and the potentially functional regions of the viral genome (Figure 1B). 60 well-folded structures were further identified in these regions, which exhibited consistent conformations in two replicates (Figure 2—figure supplements 2-11). The low SHAPE-high Shannon regions (11.70%), which exhibit variable conformations, might perform different functions through conformational transitions (Figure 1C). Designing drugs targeting these regions would require identifying the specific conformations that are responsible for the functions.

Regions with low base-pairing rates (high SHAPE) accounted for 36.57% of the PEDV genome, including the TRS-Ls of the 5’ UTRs (positions 57-72 nt) and FSE regions (positions 12605- 12690 nt) (Figure 1D-E). RNAs in the high SHAPE-high Shannon regions (26.90%) are prone to conformational shifts, and their high accessibility suggests that they are more susceptible to interact with small molecule drugs (Figure 1D). In contrast, RNAs in the high SHAPE-low Shannon regions (9.67% of the PEDV genome) maintain a stable single-stranded state within the cell, facilitating pairing with exogenous complementary sequences (Figure 1E). Consequently, these regions are considered to be ideal targets for designing antiviral oligonucleotides (Manfredonia et al., 2020; Piasecka et al., 2020; Sagan et al., 2010).

### Not all folded functional structures are potential antiviral targets

Although well-folded structures are thought to be potential targets for antiviral molecules(Childs-Disney et al., 2022; Huston et al., 2021; Manfredonia et al., 2020; Sreeramulu et al., 2021), there have been relatively limited experimental validations. The 5’UTR-SL5 of coronaviruses is required for defective interfering (DI) RNA replication(Brown et al., 2007) and important for viral packaging(Masters, 2019). In the PEDV genome, the 5’UTR-SL5 with four-way junction structure exhibits well-folded characteristics, similar to that of SARS-CoV-2 (Figure 2—figure supplement 12). Seven compounds (Table S4, compounds 1-7), previously reported to interact with the 5’ UTR-SL5 of SARS-CoV-2 (Sreeramulu et al., 2021), were found to bind with the 5’ UTR-SL5 of PEDV *in vitro*, and two compounds (compounds 1 and 4) exhibited dissociation constants (*K_D_)* at the µM-level (Figure 2—figure supplement 13). However, none of these compounds inhibited the proliferation of PEDV at 50 µM in cells (Figure 2—figure supplement 14). This result suggests that compounds bound to well-folded functional structures *in vitro* do not necessarily exhibit antiviral activity in cells. The accessibility of compounds to potential RNA targets could be an important issue, suggesting that the targetability of the 60 well-folded RNA structures in the PEDV genome requires further validation.

### Identification of PQSs in dynamic single-stranded regions

Dynamic RNAs in the high SHAPE-high Shannon regions can be induced by compounds to form stable structures that perform specific biological functions(Ratni et al., 2021). We identified 49 regions with high SHAPE reactivity and high Shannon entropy with at least 25 nucleotides in length (Table S3), spanning a total of 1748 bases (∼6.2% of the PEDV genome).

To investigate whether RNA motifs in high SHAPE-high Shannon regions can be stabilized to inhibit viral proliferation, we focused on the G4 structure in these regions. Nine putative PQSs were identified by at least three of the G4 prediction tools (Figure 3A, Figure 3—figure supplement 1). Given the abundance of PEDV mutant strains in nature, further conservation analyses identified seven PQSs that are highly conserved (> 90%) among 761 PEDV strains as potential functional G4 candidates (PQS1-PQS7, Table S5). PQS2, PQS3, PQS4, and PQS6 form stable double-stranded structures with low SHAPE reactivity and low Shannon entropy, making them less likely to fold into stable G4 structures (**Figure 3—figure supplements 2-5**). PQS5 and PQS7 did not fall in our four defined regions and were also omitted from further investigation. Conversely, PQS1 is located in the dynamic single-stranded region with high SHAPE reactivity and high Shannon entropy (Figure 3B-C), with relatively little competition from local hairpin structure folding.

**Figure 3.**
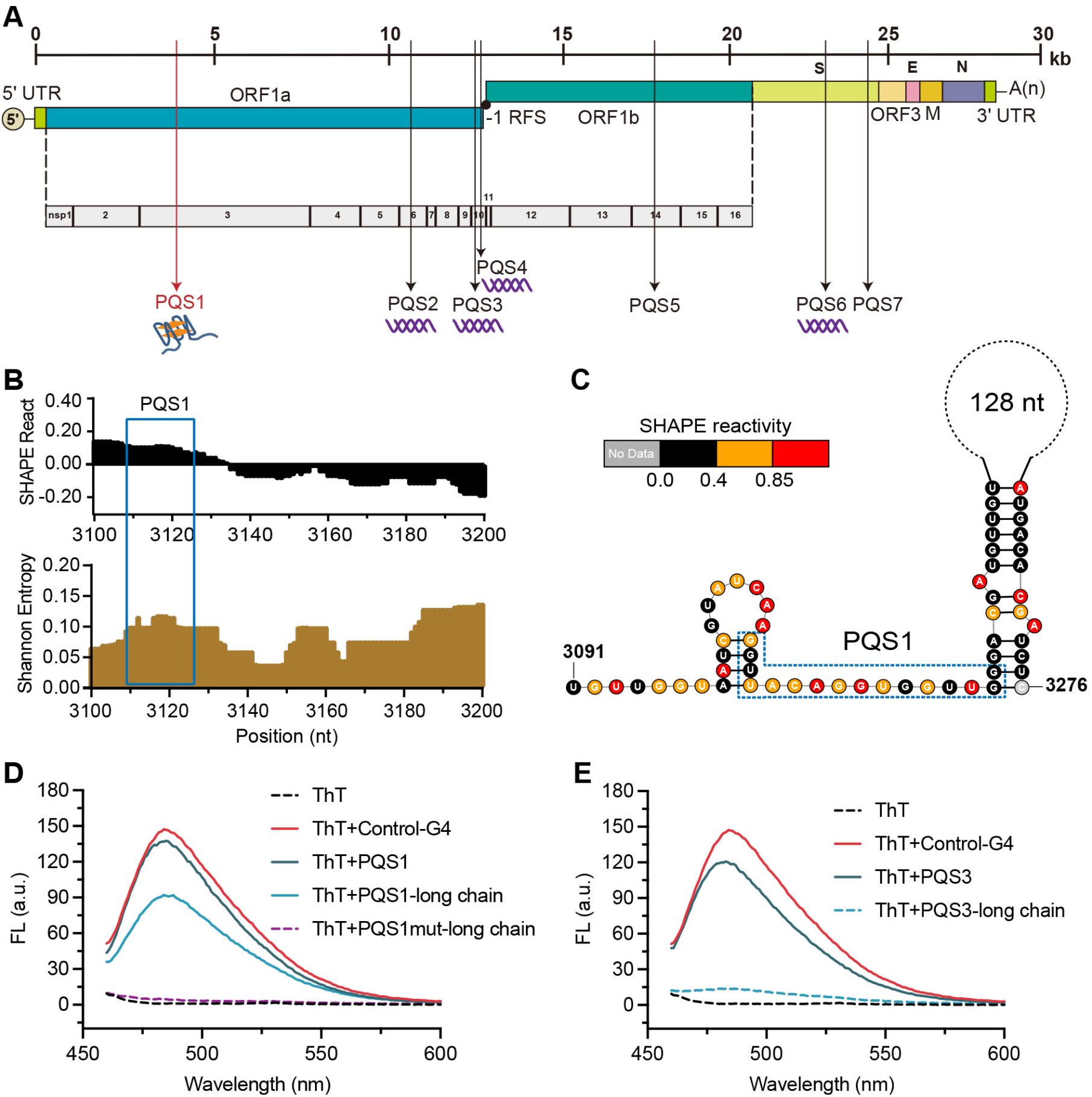
The PQS1 within high SHAPE-high Shannon regions has the potential to form G4 structure in the PEDV genome. **(A)** Distribution of PQSs in the PEDV genome. **(B)** SHAPE reactivity and Shannon entropy for the regions containing PQS1. **(C)** Local secondary structure of the region containing PQS1 predicted with SHAPE reactivity constraints. PQS1 is marked with a blue dashed box. **(D)** Fluorescence turn-on assays of ThT (1 μM) in the presence of PQS1 (0.5 μM), PQS1-long chain (0.5 μM), and PQS1mut-long chain (0.5 μM). **(E)** Fluorescence turn-on assays of ThT (1 μM) in the presence of PQS3 (0.5 μM) and PQS3-long chain (0.5 μM). Excitation wavelength was 442 nm. PRRSV-G4 RNA was used as control.

To validate whether SHAPE analysis could reflect the competitive conformational folding of PQSs in the PEDV genome, we performed *in vitro* transcription to obtain local intact structures containing PQSs within dynamic single-stranded regions and stable double-stranded regions (Table S6). Thioflavin T (ThT) fluorescence turn-on assays were conducted under physiological K⁺ conditions (100 mM), with the G4 sequence of porcine reproductive and respiratory syndrome virus (PRRSV) serving as a positive control (Control-G4)(Fang et al., 2023). The results demonstrated that for short PQSs sequences containing only G4-forming motifs (Table S7), PQS1, PQS3, PQS4, and PQS6 all induced significant ThT fluorescence enhancement (Figure 3D-E, Figure 3—figure supplement 6), confirming their ability to form G4 structures. However, in long RNA fragments encompassing PQSs and their flanking sequences, only PQS1 and PQS4 exhibited pronounced ThT fluorescence responses (Figure 3D-E), whereas PQS2, PQS3, and PQS6 showed negligible signals (Figure 3E, Figure 3— figure supplement 6). Notably, the PQS1-long chain displayed the strongest fluorescence signal, while its mutant counterpart (PQS1mut-long chain) exhibited the lowest background fluorescence (Figure 3D). These findings indicate that although most PQSs can form G4 structures *in vitro*, PQS1—located in the high SHAPE-high Shannon entropy region— demonstrates the most robust G4-forming capability when competing with local secondary structures in the genomic context. Therefore, PQS1 was selected for further structural and functional validation.

PQS1 is located in the nsp3 gene of the PEDV genome (Figure 3F) and is highly conserved in all 761 PEDV strains (98.37%, Figure 3—figure supplement 7A). Native polyacrylamide gel electrophoresis (PAGE) and circular dichroism (CD) measurements were conducted to further verify the formation of the PQS1 G4 structure. The migration rate of PQS1 noticeably accelerated with increasing K^+^ concentration (Figure 3—figure supplement 7B), whereas the migration rate of the mutant sequences (PQS1m) that did not form G4 remained relatively constant. CD spectra revealed that PQS1 adopted a parallel G4 structure in the presence of K^+^, with the corresponding typical features of positive CD band at approximately 265 nm and a negative peak near 240 nm (Figure 3—figure supplement 7C). ^1^H nuclear magnetic resonance (NMR) was utilized to further characterize the G4 structure of PQS1. Chemical shifts in the 10-12 ppm range are considered a hallmark of G4 structures(Adrian et al., 2012). The ^1^H NMR spectrum of PQS1 displayed prominent imino proton peaks in this region, confirming the formation of the PQS1 G4 structure (Figure 4A).

**Figure 4.**
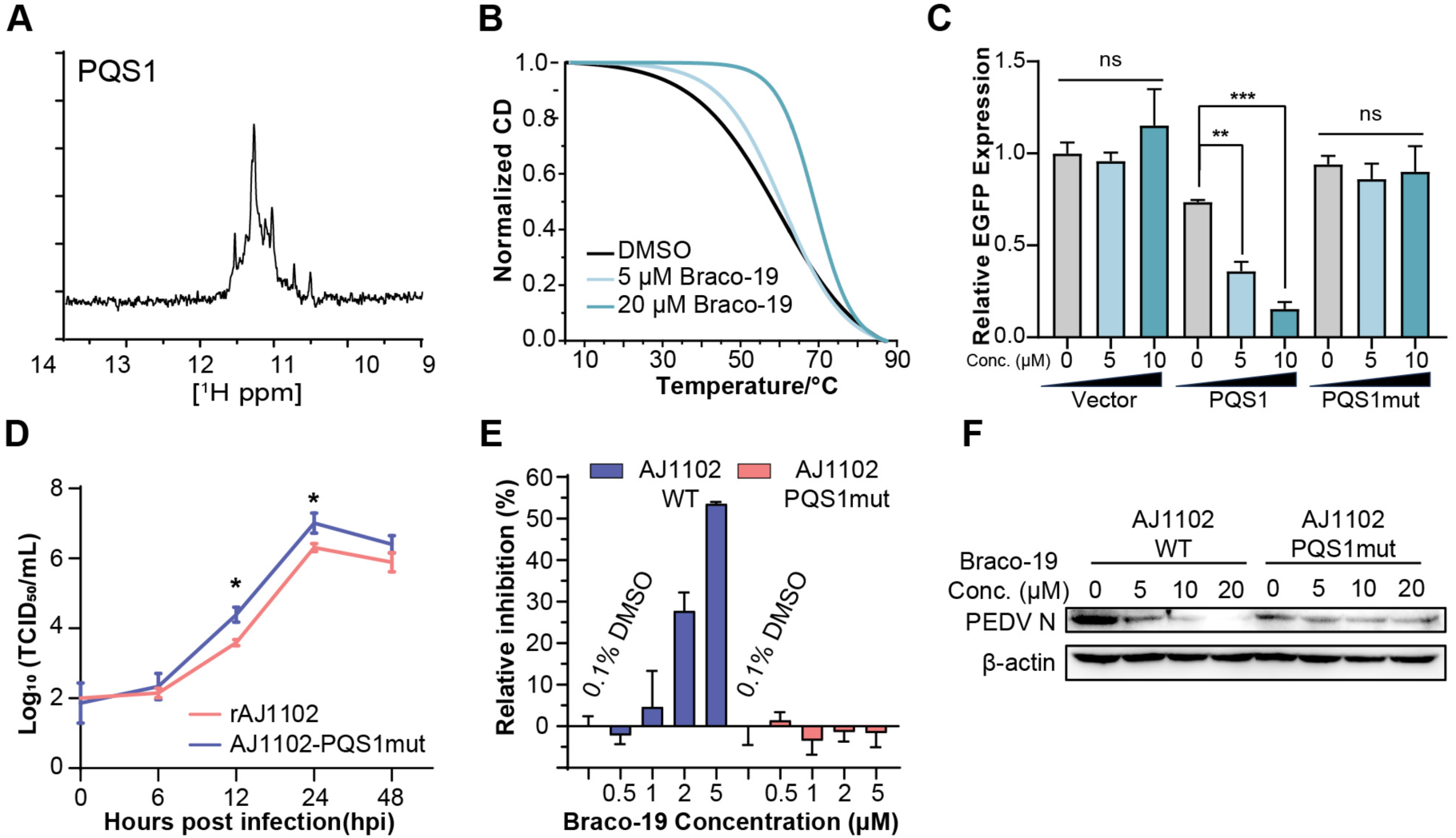
The G-quadruplex structure, biological functions of PQS1, and antiviral effects of Braco-19. **(A)** ^1^H NMR analysis of PQS1. **(B)** CD melting profiles of PQS1. **(C)** Quantitative fluorescence signal using the corrected total cell fluorescence method for EGFP in cells transfected with plasmids containing the empty vector, PQS1 and PQS1mut. **(D)** Proliferation curve of the PEDV wild type (WT) strain and PQS1 mutant strain. (**E**) The relative inhibition rates of Braco-19 against AJ1102-WT and AJ1102-PQS1mut. (**F**) Western blot analysis of the effects of Braco-19 on the viral N protein expression of AJ1102-WT and AJ1102-PQS1mut.

### Stabilization of PQS1 as a G-quadruplex inhibits RNA synthesis, gene expression and viral proliferation

The addition of the G4-specific ligand Braco-19 significantly increased the melting temperature (Tm) of PQS1 by 10.8 ℃ (Figure 4B), indicating that the RNA G4 structure of PQS1 is stabilized by Braco-19. Furthermore, the RNA stop assays and the EGFP expression inhibition experiments highlighted the effect of a stable G4 structure of PQS1 in inhibiting RNA replication (Figure 4—figure supplement 1) and protein expression (Figure 4C, Figure 4—figure supplement 2).

To explore whether PQS1 could be a potential drug target, we constructed a recombinant virus without PQS1 (AJ1102-PQS1mut) through a synonymous mutation (G3109A) that does not change the amino acid code (Figure 4—figure supplement 3A). The proliferation level of AJ1102-PQS1mut was significantly faster than that of the wild-type strain rAJ1102, suggesting that disruption of the folding of the PQS1 facilitates viral proliferation (Figure 4D). Another potential G4 sequence, PQS3, located in a stable double-stranded region with low SHAPE reactivity and low Shannon entropy, was used as a control. The PQS3 mutant recombinant virus (AJ1102-PQS3mut) proliferated at the same level as the wild type after synonymous mutation (G12322A) of its G4 sequence (Figure 4—figure supplement 3B).

Braco-19 inhibited PEDV proliferation with a median effective inhibitory concentration (EC50) of 3.60 µM (Figure 4—figure supplement 4A), which is well below the median cytotoxic concentration (CC50 > 100 µM) of Braco-19 (Figure 4—figure supplement 4B). In contrast, no inhibition was observed for Braco-19 against the mutant strain AJ1102-PQS1mut (Figure 4E). Western blot analysis was used to determine the effects of Braco-19 on viral protein expression in infected cells using the PEDV N protein antibody. The production of AJ1102-WT N protein was significantly reduced with increasing concentrations of Braco-19, while no change was observed in the production of AJ1102-PQS1mut N protein (Figure 4F), indicating that Braco-19 inhibited protein expression of PEDV in cells by targeting PQS1. Furthermore, as a control, we observed nearly identical inhibitory activity of Braco-19 against both the PQS3 mutant strain (AJ1102-PQS3mut) and wild-type virus (Figure 4—figure supplement 3C), demonstrating the specificity of Braco-19’s action on PQS1.

### Efficient antiviral siRNAs design targeting stable single-stranded regions

High SHAPE-low Shannon regions represent stable, accessible RNAs in cells, likely exposing long Watson–Crick edges of multiple residues available for base-pairing with complementary sequences (Figure 1D). Our analysis of fold characterisation (see Methods) identified 73 such regions (Table S3) with at least 25 nucleotides in length, spanning a total of 2696 bases (∼9.6% of the PEDV genome). To test the suitability of these regions as targets for antiviral siRNAs, we designed four siRNAs targeting these regions that (i) have at least 10 consecutive unpaired nucleotides, and (ii) were highly conserved across the different PEDV strains (Table S7, Figure 5A-D). As a control, we designed four additional highly conserved siRNAs targeting duplex regions (Figure 5—figure supplement 1) and one non-targeting siRNA that does not target any sequence of the viral and host genomes (Table S7).

**Figure 5.**
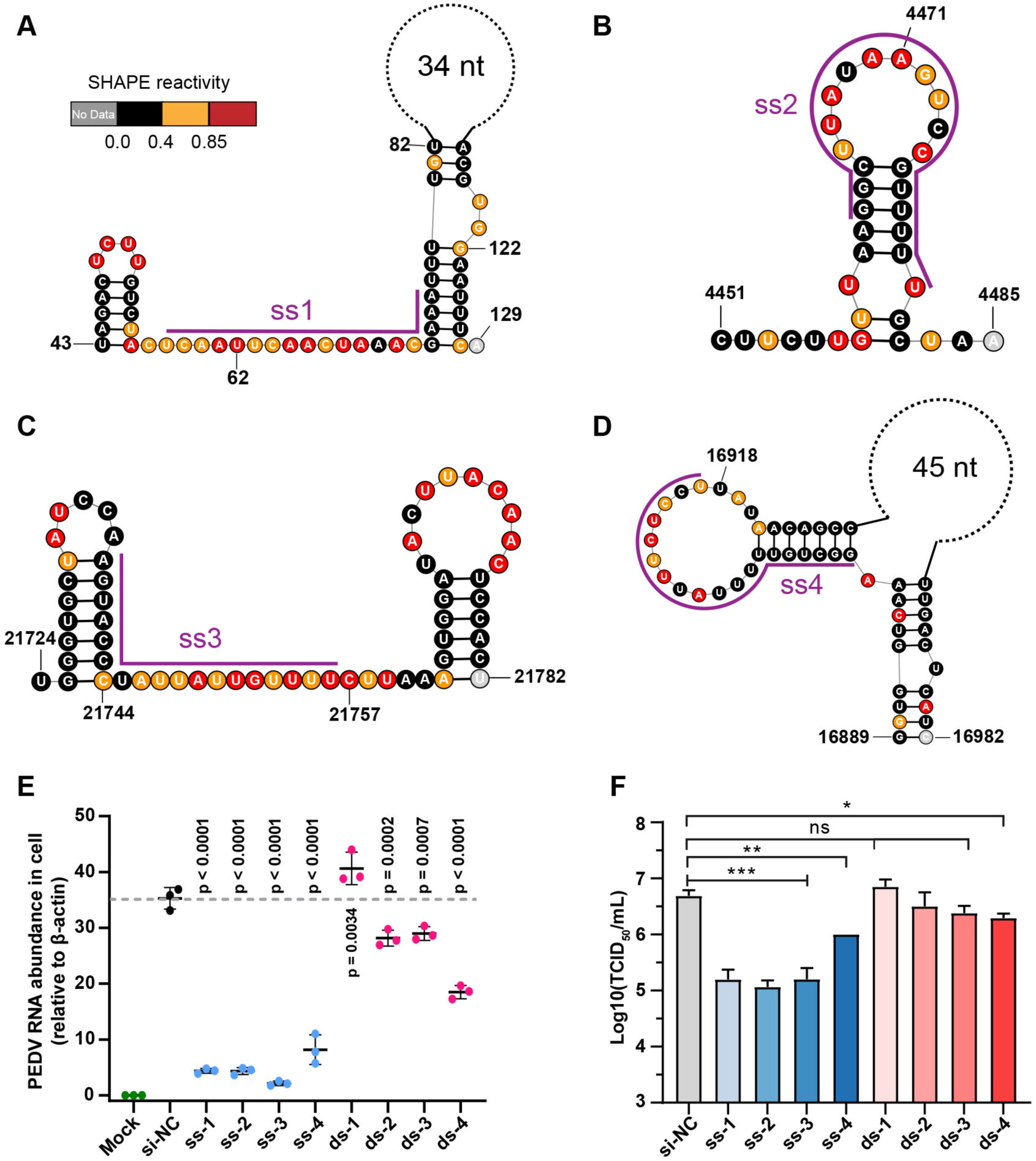
Secondary structure of target regions and antiviral effects of siRNAs. **(A-D)** Local secondary structures of the stable single-stranded regions targeted by siRNAs predicted by SHAPE reactivity as a constraint; (A) ss1; (B) ss2; (C) ss3; (D) ss4. **(E)** qPCR showing the relative abundance of the PEDV RNA genome in infected Vero cells. The four siRNAs targeting the high SHAPE-low Shannon regions and the four siRNAs targeting the duplex regions are labeled ss-1 to ss-4 and ds-1 to ds-4, respectively. ss (single-stranded targeting siRNAs); ds (dual-stranded targeting siRNAs). si-NC was a control siRNA that did not target any viral or host sequences, and the mock group was not inoculated with virus. **(F)** TCID50 assays for detecting virus titers. The presented results represent the means and standard deviations of data from three independent experiments. ns: no significant difference. * P < 0.05; ** P < 0.01; *** P < 0.001, Duncan’s multiple comparison test.

RT‒qPCR results of infected cells at 16 h post viral exposure revealed that, four siRNAs targeting high SHAPE-low Shannon regions significantly inhibited viral proliferation in cells (Figure 5E) compared with the non-targeting siRNA control (siRNA-NC). Consistent with the RT-qPCR results, compared with the siRNA-NC, we observed that these four siRNAs had minimal cytopathic effects (CPEs) in Vero cells (Figure 5—figure supplement 2). Further TCID_50_ analysis demonstrated that, three out of the four tested siRNAs targeting single-stranded regions reduced the TCID_50_ by approximately 1.5log10 titer compared to the siRNA-NC (ss-1, ss-2, and ss-3; Figure 5F). In contrast, only one of the four siRNAs targeting duplexes could slightly prevent proliferation of PEDV in cells (ds4; Figure 5E-F, Figure 5—figure supplement 2). The target sequences of four single-stranded targeting siRNAs are more than 85% conserved across 761 known PEDV strains. Combining two or more of these siRNAs is expected to be effective in inhibiting nearly all known PEDV strains. These results showed that siRNAs targeting single-stranded regions with high SHAPE reactivity and low Shannon entropy can be more effective at exerting their antiviral effects.

## DISCUSSION AND CONCLUSIONS

Several RNA-targeting antiviral strategies have highlighted the potential of therapeutic application(Dai et al., 2024). However, it is challenging to identify these RNAs from the viral genome *in situ*. In SHAPE-MaP analysis, regions with low reactivity and low entropy values are more likely to form a single stable structure in cells, and are therefore considered candidates for potential functional RNAs and targets for antiviral therapy. Novel well-defined RNA structures that are critical for viral fitness in HIV, HCV, DENV and SINV have been identified by the analysis of low SHAPE and low entropy regions (Dethoff et al., 2018; Kutchko et al., 2018; Mauger et al., 2015; Siegfried et al., 2014). However, there are still no clinically approved small molecule drugs or oligonucleotides targeting stable folded RNA structures. Our data shows that two compounds that interacted with the well-folded 5’UTR-SL5 of PEDV *in vitro* at µM-level *K_D_*, but did not show antiviral activity at 50 µM in cells (Figure 2—figure supplements 13-14). This result suggests that, on the one hand, the low accessibility of the regions with low SHAPE reactivity makes them difficult to access by compounds in cells, and may require compounds to have better *K_D_* to RNA. On the other hand, the binding of compounds to the well-folded structure does not necessarily alter its function. In contrast, the FDA-approved compound Risdiplam exerts its biological activity by stabilizing transient RNA structures(Campagne et al., 2019; Ratni et al., 2021; Sheridan, 2021), suggesting that it may be possible to inhibit viral proliferation by stabilizing variable conformations. We therefore focused our attention on highly accessible regions with high SHAPE reactivity in cells.

We identified 42 low SHAPE and high Shannon regions and 49 high SHAPE and high Shannon regions in the PEDV genome by SHAPE-MaP, representing conformationally variable RNAs in the viral genome. As the low SHAPE regions are highly structured, it is difficult for the G4 in these regions to outcompete the folding of long base paired structures (Figure 6A). We therefore selected a potential G4 sequence (PQS1) located in the high SHAPE-high Shannon region of nsp3 (PLpro), where a G4-disrupting mutation promotes viral proliferation. We found that the compound Braco-19 significantly stabilised the G4 structure of PQS1 *in vitro* and inhibited PEDV proliferation in cells. Crucially, Braco-19 showed no inhibitory activity against the PQS1-mutant strain while maintaining potent activity against the PQS3-mutant strain (Figure 4E, Figure 4—figure supplement 3C). This suggests that the compound can selectively target the PQS1 of the high SHAPE-high Shannon region in cells. This approach can be extended to other RNA viruses and potentially to DNA viruses where RNA intermediates play critical roles in the viral life cycle.

**Figure 6.**
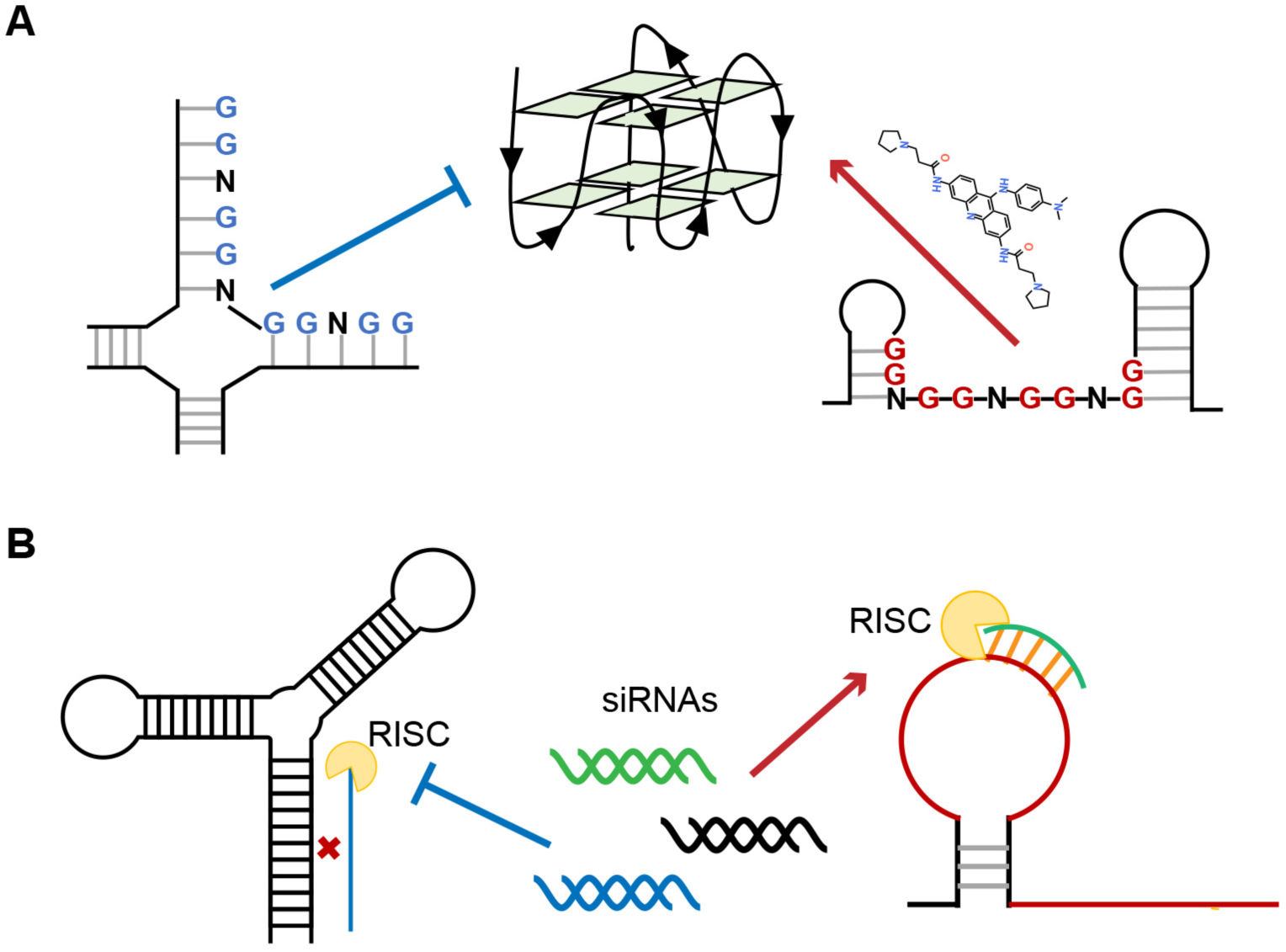
Single-stranded RNA regions in viral genomes as the targets for antiviral therapy. **(A)** PQSs in the single strands are easier to be induced into G4s by ligands than those in the paired regions. (**B**) Influence of the secondary structures on the binding efficiency of siRNAs.

Recently, the FDA has approved several siRNAs for therapeutic purposes(Ali Zaidi et al., 2023). siRNA-mediated RNAi is particularly suited to inhibit viruses with single-stranded RNA genomes due to its potential to specifically silence virtually any therapeutic target. When screening siRNA target sequences, sequence conservation and the functional importance of the target region are usually given priority. However, the RNA structure of the target region also has a significant impact on the efficiency of RNAi (Figure 6B)(Cao et al., 2021; Manfredonia et al., 2020; Patzel et al., 2005; Piasecka et al., 2020; Sagan et al., 2010; Szabat et al., 2020). Our approach uniquely integrates in situ RNA structural data (SHAPE reactivity and Shannon entropy) to prioritize siRNA targets within stable single-stranded regions (high SHAPE reactivity, low Shannon entropy), which are experimentally validated as accessible in infected cells. This represents a significant departure from traditional siRNA design methods that rely primarily on sequence conservation, thermodynamic rules (e.g., Tuschl rules), or *in vitro* structural predictions(Ali Zaidi et al., 2023; Qureshi et al., 2018; Tang and Khvorova, 2024),which may not accurately reflect intracellular RNA accessibility. Bowden-Reid et al. designed 39 antiviral siRNAs against various SARS-CoV-2 variants based on sequence conservation, ultimately identifying 8 highly effective sequences(Bowden-Reid et al., 2023). Notably, five of these effective sequences targeted regions that were located in high SHAPE-high Shannon regions according to SARS-CoV-2 SHAPE datasets (Table S8)(Manfredonia et al., 2020). This independent finding aligns perfectly with our conclusions and demonstrates that SHAPE-based siRNA design outperforms sequence/structure-agnostic approaches, at least in terms of significantly improving antiviral siRNA screening efficiency. Given the growing availability of SHAPE datasets for numerous viruses, we are confident that our methodology will facilitate more precise design of antiviral siRNAs.

## METHODS

### Cell culture and PEDV infection

Vero E6 cells were cultured in 10 cm culture dishes in Dulbecco’s modified Eagle’s medium (DMEM) supplemented with 10% fetal bovine serum (FBS) at 37°C in an atmosphere of 5% CO_2_. The cells were infected at an MOI of 0.1 with PEDV/AJ1102 (GenBank accession: MK584552.1) when the cell confluence reached 95% or greater.

### *In situ* RNA probing and purification

During the logarithmic phase of viral genome replication after infection (12 h), infected cells were washed 3 times with 10 ml cold PBS and then resuspended in PBS containing 2-methylnicotinic acid imidazolide (NAI(Spitale et al., 2013)) at a final concentration of 100 mM. Two independent replicates were performed for *ex vivo* experiments. An equal amount of DMSO was added to another sample as a control. The samples were then incubated at room temperature with continuous shaking for 15 minutes, after which the reaction was terminated by the addition of DTT at a final concentration of 0.5 M. The supernatant was then removed, and the cells were lysed by adding 6 ml of TRIzol. The RNA was extracted by adding 1.2 ml of chloroform:isoamyl alcohol (24:1). Three milliliters of the upper aqueous phase was transferred to a new tube, 6 ml of anhydrous ethanol was added, and the mixture was incubated overnight at -20°C. The RNA was pelleted at 18,000 × g for 20 min at 4°C, washed twice with 80% EtOH, and then the pellet was air-dried. Finally, RNA was dissolved in RNase-free water and then stored at -80°C until further use.

### Isolation and fragmentation of poly(A)PEDV genomic RNA

We isolated poly(A) RNA with VAHTS mRNA capture beads (Vazyme #N401) according to the manufacturer’s instructions. Five micrograms of total RNA was diluted to 50 μl with RNase-free water and purified in two rounds with mRNA capture beads. Poly(A) RNA was fragmented to a median size of 150-200 nt by incubation at 94°C for 8 min in fragmentation buffer (65 mM Tris–HCl; 4 mM MgCl_2_; 95 mM KCl; pH 8.0) and then purified with an RNA Clean & Concentrator-5 kit (Zymo Research, R1015) following the manufacturer’s instructions.

### Reverse transcription of modified PEDV genomic RNAs

20 μl of poly(A) RNA (∼500 ng) was added to separate 0.65 ml RNase-free tubes, 2 µl of 200 ng/µl random primer was added to each tube, and the mixture was incubated at 65°C for 5 min. Sixteen microliters of 2.5× MaP buffer (125 mM Tris, pH 8.0; 187.5 mM KCl; 15 mM MnCl_2_; 25 mM DTT and 1.25 mM dNTPs) were added to each tube, and the mixture was incubated at 25°C for 2 min. Then, 2 μl of SuperScript II reverse transcriptase (200 U/µl; Life Technologies, cat. no. 18064-014) was added, and the mixture was incubated at 42°C for 3 h. The reactions were heat-inactivated by incubation at 75°C for 15 min, after which the mixture was placed on ice. cDNA was purified from the reactions using G-25 micro spin columns (GE Healthcare, cat. no. 27-5325-01) following the manufacturer’s instructions. The second-strand cDNA synthesis reaction was then performed immediately. The volume of 1st strand cDNA was adjusted to 56 μl with nuclease-free H_2_O, 6.5 μl of NEBNext Second Strand Synthesis Reaction Buffer (10×; New England Biolabs, cat. no. E6111S) and 3.5 μl of NEBNext Second Strand Synthesis Enzyme Mix (New England Biolabs, cat. no. E6111S) were added, and the mixture was then incubated at 16°C for 2.5 h.

### SHAPE-MaP sequencing library construction and quality control

Sequencing libraries were generated using a VAHTS^®^ Universal V8 RNA-seq Library Prep Kit for MGI (Vazyme #NRM605) according to the manufacturer’s protocol. Libraries concentrations were measured using a Qubit dsDNA HS Assay Kit (Thermo Fisher, Cat. No. Q32851). One microliter of the final library product was removed, and the library quality was analyzed using an Agilent DNA1000 Kit (Agilent, Cat. No. 5067-1504).

### SHAPE-MaP data analysis

SHAPE-MaP data analysis was adapted from Manfredonia *et al*^13^, and performed using the RNA Framework v2.6.9(Incarnato et al., 2018) (https://github.com/dincarnato/RNAFramework), employing the following steps - a) Data was preprocessed and aligned to the PEDV reference genome using the rf-map module with the parameters: -ctn -cmn 0 -cqo -cq5 20 -b2. Reads with internal Ns were discarded, and low-quality bases (Phred < 20) were excluded before alignment. b) The index of the PEDV genome was built using Bowtie2 software (Langmead and Salzberg, 2012). Individual nucleotide mutation signals were assessed with the rf-count module with parameters: -m -rd, generating a binary RC file with sequence data, mutation counts, coverage, and total aligned fragments. c) The rf-norm module was applied for data normalization with parameters: -sm 3 -nm 3 -rb AC -n 500 -mm 1, utilizing the scoring method developed by Siegfried *et al*(Siegfried et al., 2014).

### PEDV RNA secondary structure modeling guided by SHAPE data

The rf-fold module, employing a previously established windowed approach^13,^ (Siegfried et al., 2014), predicts the RNA secondary structure of the entire PEDV genome using the normalization step’s XML file, with a default pseudo-free energy intercept and slope values of -0.6 (kcal/mol) and 1.8 (kcal/mol), respectively, with the following parameters: -fw 3000 -fo 300 -wt 200 -pw 1000 -po 250 -dp -sh -nlp -md 600. This means that RNAstructure will generate a 3000 nt window and slide in 300 nt increments over the genome in 1000 nt partition windows, predicting the most reasonable secondary structure of the genome under the condition that the maximum base pairing distance does not exceed 600 nt. SHAPE reactivities were incorporated in the form of soft constraints using the Vienna RNA package (Lorenz et al., 2016). For each window, the first and last 100 nt were ignored to avoid terminal biases. Additionally, RF Fold can provide pairing probabilities between bases, Shannon entropy of individual bases, and potential pseudoknot structures. The Shannon entropy is commonly used to measure the diversity of RNA secondary structures:

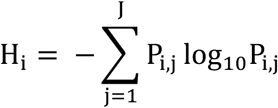

where P represents the pairing probability between bases.

### Identification of characterized regions in viral genomes

Based on the previous research methods for identifying well-folded regions ^13,^ (Siegfried et al., 2014), ^25, 29^, modifications have been made as follows:

**a)** For the identification of low SHAPE-low Shannon regions, we first calculated the local medians of SHAPE reactivities and Shannon entropies in sliding, centered, 50 nt windows and subtracted the global median for visualization. Then, a window of 50 nt was slid along the genome with a 1 nt step. Windows with >75% of the bases below both the global SHAPE and Shannon median (calculated on the full PEDV genome) were selected, and windows less than 10 nt apart were merged. Following the initial identification of well-folded regions, entire structures exhibiting consistent conformations in both datasets and having at least 50% of their bases in well-folded regions were further selected. Subsequently, well-folded RNA structures within cells were determined.
**b)** For the identification of low SHAPE-high Shannon regions, a window of 50 nt was slid along the PEDV genome. Windows with >75% of the bases below the global SHAPE median and above the global Shannon median (based on the full PEDV genome) were selected, and windows less than 10 nt apart were merged.
**c)** For the identification of high SHAPE-low Shannon regions, instead, a window of 25 nt was slid along the PEDV genome. Windows with >75% of the bases were above the global SHAPE median and below the global Shannon median (calculated on the full PEDV genome), and >50% of the bases were predicted to be single-stranded in the MEA structure; windows less than 10 nt apart were selected and merged.
**d)** For the identification of high SHAPE-high Shannon regions, a window of 25 nt was slid along the PEDV genome. Windows with >75% of the bases above both the global SHAPE and Shannon median (based on the full PEDV genome) were selected, and windows less than 10 nt apart were merged.

### G-quadruplex forming sequence prediction and conservativeness analysis

The G4 sequence was defined as G_≥2_N_1-6_G_≥2_N_1-6_G_≥2_N_1-6_G_≥2_, where G refers to guanine and N refers to any nucleotide, including guanine. Four independent G4 prediction tools (QGRS- mapper, G4Catchall, ImGQfinder and pqsfinder) were employed to analyse the potential G4 forming sequences (PQSs) in the PEDV genome(Doluca, 2019; Hon et al., 2017; Kikin et al., 2006; Varizhuk et al., 2017). Conservation analysis was performed using Molecular Evolutionary Genetics Analysis (MEGA) software (version 6.0; available at www.megasoftware.net). 761 PEDV genome sequences were retrieved from the National Center for Biotechnology Information website (NCBI, https://www.ncbi.nlm.nih.gov/). The graphical representation of the alignment sequence was generated by WebLogo 2 (http://weblogo.berkeley.edu)(Crooks et al., 2004).

### Verification of G-quadruplex structure formation by native PAGE

Cy2-tagged RNA was annealed in buffers (100 mM potassium arsenate, pH 7.0) containing different concentrations of KCl (0, 50 mM, 100 mM). The marker used in this experiment was a mixture of P15 (UAAUACGACUCACUA), M17 (UAAUACGACUCACUAUAUA), and M20 (UAAUACGACUCACUAUACGA). Acrylamide at a 20% concentration was used to separate the gel. The concentration of the RNA sample was 100 nM, and electrophoresis was conducted at 100 V for 2 h. Native PAGE was carried out under a controlled temperature (4.0°C) with a vertical electrophoresis instrument (DYCZ-22A; Bio-Rad, Beijing, China). After electrophoresis, RNA oligomers were scanned by a Pharos FX molecular imager for visualization.

### Circular dichroism (CD) spectroscopy

RNAs (20 μM) were dissolved in 10 mM Tris-HCl buffer (pH 7.0) containing varying concentrations of KCl (0, 50, 100 mM). The CD experiment was performed at 25°C using a JASCO-810 spectrophotometer (Jasco, Easton, MD, USA) with a quartz cell path length of 1.0 mm. CD spectra were collected from 320 to 200 nm with a scanning speed of 50 nm/min. All CD spectra were baseline corrected for the signal contribution of the buffer and represent the mean of at least two repeats.

### Nuclear magnetic resonance (NMR) spectroscopy

^1^H NMR spectra were recorded at 298 K using an 800-MHz Brock Avance DRX-800 spectrometer equipped with a low-temperature triple-resonance reverse autotuning and matching probe. The presumption of water is achieved by excitation engraving. The RNA samples were dissolved in DEPC solution containing 200 mM KCl and 10% D_2_O at a final concentration of 0.5 mM. The sample was heated at 95°C for 5.0 min and cooled slowly to 25°C. The facility was set for 100 ms diffusion time (TD), 50 ms eddy recovery time (TE), and 2 SEC relaxation delay (TR).

### CD melting studies

The compound Braco-19 was dissolved in DMSO at a concentration of 20 mM. RNA samples were dissolved in buffer containing 10 mM Tris-HCl (pH 7.0) and 100 mM KCl at a concentration of 20 mM. The samples were heated at 95°C for 10 min and then slowly annealed to room temperature. CD melting was measured on a Jasco-810 spectrophotometer equipped with a water bath temperature control attachment. CD melting curves were recorded using a heating rate of 0.2°C/min, and absorbance values were collected at 1°C intervals. The thermal stability of G4 RNA was measured by recording the CD value at 264 nm in relation to temperature. The required temperature range was set from 4.0 ℃ to 98 ℃.

### RNA stop assay

P15 (300 nM) and template RNA (600 nM) were annealed in buffer at 25°C [50 mM HEPES, 20 mM NaCl, 1.0 mM KCl, 5 mM MgCl_2_ and 4 mM DTT (pH 7.0)], heated at 95°C for 5.0 min and slowly cooled to 4°C. Subsequently, NTPs (final concentration, 200 μM) and 3Dpol (recombinant RNA-dependent RNA polymerase; Abcam, ab277617, 0.02 mg/reaction) were added to the system, and the reaction was carried out at 33°C for 20 min. An equal amount of stop buffer (95% formamide) was added, and the reaction was stopped after heating at 90°C for 4.0 min. The products were loaded and separated on a 20% denatured polyacrylamide gel. Finally, the gel was scanned using a Pharos FX molecular Imager (Bio-Rad) operated in fluorescence mode.

### EGFP expression repression by RNA G-quadruplex stabilization

We constructed pEGFP-C1 vector with wild-type PQS1 sequences or G4-mutated PQS1mut sequences fused to the N terminus of an EGFP reporter gene. Vero cells were transfected with Lipofectamine 3000 (Invitrogen, USA) according to the manufacturer’s protocol and incubated with media supplemented with various concentrations of Braco-19(Gowan et al., 2002) for 48 hours. The cells were then washed with PBS and fixed at 25°C for 15 minutes with 4% paraformaldehyde. The samples were washed three times with PBS for 5 minutes each and then fixed with precooled methanol at -20°C for 10 minutes. Observations were conducted using a two-photon laser scanning confocal microscope (Olympus, fv1000mp).

### Construction of the RNA G4-mutated recombinant virus

The pBAC-CMV-PEDV template was obtained from Prof. Xiao Shaobo (College of Veterinary Medicine, Huazhong Agricultural University). The primers used were chemically synthesized (Table S1). The sequence of the G-rich region (3109-3125 nt) was mutated (AAGGTTACAGGTGGTTGG to AAAGTTACAGGTGGTTGG). Two pairs of primers (PQS1Mut-F1 and PQS1Mut-R1, PQS1Mut-F2 and PQS1Mut-R2) were used to amplify the fragment containing the mutant site. The PQS1mut fragment containing mutation sites was amplified through overlap PCR. A scaffold oligo was used as a template, PQS1m-sgRNA-a and PQS1m-sgRNA-a were used as primers to synthesize guide DNA, and then, guide RNA (sgRNA) was obtained by reverse transcription. pBAC-CMV-PEDV was cut with the Cas9 enzyme and sgRNA. The PQS1mut fragment was cloned and inserted into a linearized pBAC- CMV-PEDV vector to generate the pBAC-PEDV-PQS1mut mutant plasmid. Vero cells were placed on 6-well plates, and plasmid transfection was performed when the cells grew to 60% to 80% confluence according to the manufacturer’s protocol (Lipofectamine 3000, Thermo, L3000015). If no lesions appeared, the freeze‒thaw solution was added to the six-well plate with 80% confluence after three repeated freeze‒thaw cycles for further culture. When lesions appeared, successful mutation of the mutation site was confirmed by reverse transcription sequencing, and the PEDV-PQS1mut mutant strain was obtained.

### Determination of the antiviral activity of Braco-19 against PEDV by RT‒qPCR

Braco-19 at different concentrations (0.5, 1, 2 and 5 μM) were inoculated into monolayer Vero cells and incubated for 2 h. PEDV-WT and PEDV-PQS1mut were inoculated at an MOI of 0.1 and incubated for 2 h. The DMEM was replaced with DMEM containing Braco-19, and the mixture was incubated for 16 h. cDNA was synthesized from 100 ng of total RNA per sample using the Transcriptor First Strand cDNA Synthesis Kit (Roche, Switzerland) according to the manufacturer’s instructions. Quantitative PCR (qPCR) was conducted following the Minimum Information for Publication of Quantitative Real-Time PCR Experiments (MIQE) guidelines (Bustin et al., 2009) using Taq Pro Universal SYBR qPCR Master Mix (Vazyme, Q712) according to the manufacturer’s protocol. mRNA expression was normalized to beta-actin levels with primers β-actin-F/R (Supplementary Table S1). PEDV genomic RNA was quantified at the nonstructural protein 12 (*nsp12*) gene using primers qPCR-nsp12-F/R (Supplementary Table S1).

### Western blot analysis to detect PEDV N gene expression

Protein samples were separated by SDS‒PAGE using a 12.5% polyacrylamide gel and then transferred to a polyvinylidene difluoride (PVDF) membrane (Millipore, USA). The membrane was blocked with 5% BSA in Tween-PBS buffer and incubated overnight at 4°C with primary antibodies. After three washes, it was incubated with HRP-conjugated secondary antibodies (Beyotime, China). Following three more washes, protein bands were detected using enhanced chemiluminescence (Bio-Rad, USA).

### CCK-8 assays

Vero cells in 96-well plates were exposed to different compound concentrations for 12 hours, followed by the addition of 10 μl CCK-8 reagent (Beyotime, China) and a 2-hour incubation at 37°C. Cell proliferation was evaluated by measuring the absorbance at 450 nm with a microplate reader.

### Indirect immunofluorescence assay (IFA)

Cells in 24-well plates were treated with Braco-19 and then infected with PEDV at 37°C for 2 hours. After washing with serum-free DMEM, cells were cultured with or without compounds in DMEM. Post-infection, cells were fixed, permeabilized, and washed before blocking and incubating with primary (mouse anti-PEDV N protein antibody) and Alexa Fluor 488-secondary antibody. Nuclei were DAPI-stained, and fluorescence images were captured using an Olympus IX73 microscope.

### Isothermal Titration Calorimetry (ITC)

ITC experiments were conducted on an AutoiTC100 titration calorimeter (MicroCal) at 25°C. All solutions (buffer, RNA, and ligand) were degassed prior to experimentation. A 10 μM solution of RNA 5’ UTR-SL5 was introduced into the cell, and a 250 μM compound solution was added to the rotating syringe (750 rpm). A total of 40 μL of compound was added to the RNA in 20 injections, with a 150 s interval between injections and an initial injection of 0.4 μL to account for diffusion from the syringe into the cell during equilibration. This initial injection was excluded from the data fitting. Additionally, dilution experiments were performed with buffer (100 mM HEPES (pH 8.0), 100 mM NaCl, and 10 mM MgCl_2_) in the cell, and the compound remained in the syringe. We utilized Microcal ITC analysis software to analyze the raw ITC data using the two-site binding model. We subtracted the dilution data from the raw interaction data of the compound with RNA prior to analysis. Following the fitting of these experimental data, the enthalpy change during the process was determined.

### siRNA antiviral assay

siRNAs were synthesized in GENERAL BIOL and transfected into 8 × 10^4^ Vero E6 cells in 24-well plates using 10 pmol of each siRNA and Lipofectamine™ RNAiMAX Transfection Reagent (Invitrogen, 13778075) according to the manufacturer’s protocol. The transfected cells were infected with PEDV (MOI = 0.1) after 24 h. After 16 hours of PEDV infection, minimal cytopathic effects (CPEs) on the Vero cells were observed under the microscope. To quantify RNA levels of PEDV in infected cells, the total RNAs from the infected Vero cells were extracted with TRIzol Reagent (Invitrogen, 15596026). For RT-qPCR, 100 ng of total RNAs were converted into cDNA using HiScript II 1st Strand cDNA Synthesis Kit (Vazyme, R212). The qPCR was performed with ChamQ Universal SYBR qPCR Master Mix (Vazyme, Q711) on a StepOnePlus real-time PCR machine (Applied Biosystems). The primers targeting the nucleocapsid (*N*) gene of PEDV and the *β-actin* gene of Vero cells were used for qPCR. For the 50% tissue culture infectious dose (TCID_50_) assay, monolayers of Vero cells seeded in 96-well plates were washed twice with DMEM. The supernatants of infected cells were serially diluted tenfold. Cells were then inoculated with eight replicates of the appropriate dilutions of the virus suspension. The TCID_50_ was calculated by the Reed-Muench method after culturing the cells for 48 h at 37°C in 5% CO_2_.

### Statistical analysis

The experimental data were analyzed by Student’s t test or one-way or two-way analysis of variance using GraphPad Prism 8 software. *P < 0.05 was considered to indicate statistical significance.

## DATA AVAILABILITY

SHAPE-MaP datasets generated in this study have been deposited at Zenodo.org (https://zenodo.org/doi/10.5281/zenodo.13743827) and include Gene Expression Omnibus (GSE271098). XML files with normalized SHAPE reactivities (SHAPE_react_rep1/2.react.xml); WIG files with Shannon entropies (SHAPE_react_rep1/2.react.xml) and the full secondary structure (PEDV_incell_secondary structure.ct).

## SUPPLEMENTARY INFORMATION

The supplementary information contains Tables S1-S8.

## AUTHOR CONTRIBUTIONS

**Dehua Luo:** Designed of the study, investigation, formal analysis, validation, writing; **Yingge Zheng:** performed the EGFP reporter vector construction and PQSs mutant virus rescue; **Zhiyuan Huang, Zi Wen, Yingxiang Deng:** SHAPE data processing. **Lijun Guo, Qingling Li, Yuqing Bai:** data visualisation; **Shozeb Haider:** validation, writing - review and editing; **Dengguo Wei:** conceptualization, supervision, validation, writing – review and editing.

## Acknowledgements

This work was supported by the China Postdoctoral Science Foundation (2024M761080), the Natural Science Foundation of Hubei Province (2025AFB288), the National Key R&D Plan of China (2021YFD1801102), the National Natural Science Foundation of China (22077043 and 32272491), the Fundamental Research Funds for the Central Universities (2662023PY005 to DW), the HZAU-AGIS Cooperation Fund (SZYJY2022016 to DW), the Natural Science Foundation of Hubei Province (2021CFA061 to DW, 2017CFB233 to SL), and funding from the National Key Laboratory of Agricultural Microbiology (AML2023B05 to DW). The funders had no role in the study design, data collection and analysis, decision to publish, or preparation of the manuscript.

---

**Figure 1—figure supplement 1.**
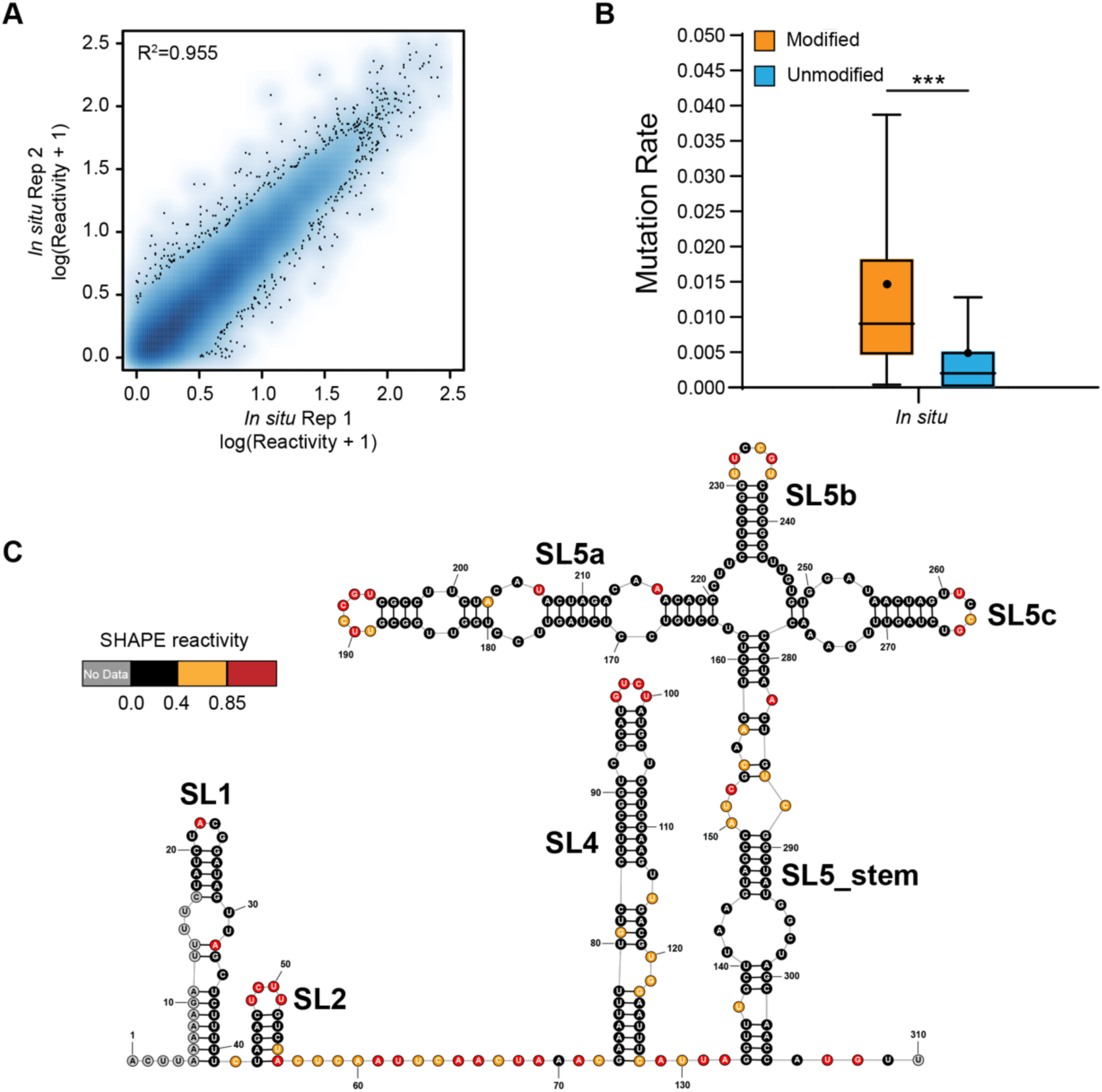
Genome-wide SHAPE-MaP analysis of PEDV. **(A)** Heat scatter plot of the SHAPE reactivities across two biological replicates for the *in vivo* datasets. **(B)** Reverse transcription mutation rates (median indicated by line, average indicated by ".") for two *in vivo* biological replicates resulting from the NAI-treated (modified) group versus the DMSO-treated (unmodified) group. A significant increase in the reverse transcription mutation rate was observed in the NAI-treated group compared to the DMSO-treated group, suggesting that NAIs are effective at modifying viral genomic RNA *in vivo*. **(C)** SHAPE secondary structure modeling of the PEDV 5’ UTR. ***p < 0.001 by equal variance unpaired Student’s t test.

**Figure 2—figure supplement 1.**
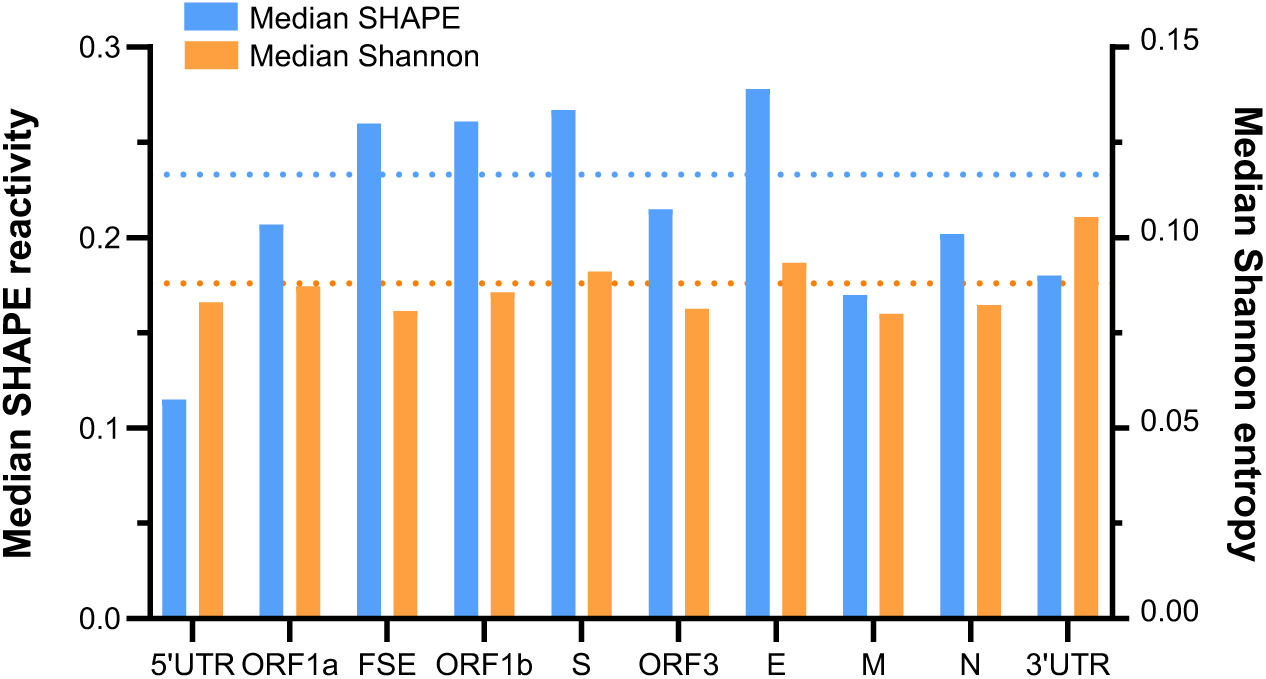
Folding characteristics of known functional regions and coding areas in the PEDV genome. The length of ORF1ab is 20,345 nt, accounting for 72.5% of the total length of the PEDV genome (28,044 nt) (a major contributor to the global median). The blue dashed line and the orange dashed line represent the global median SHAPE reactivity and the global median Shannon entropy, respectively.

**Figure 2—figure supplement 2.**
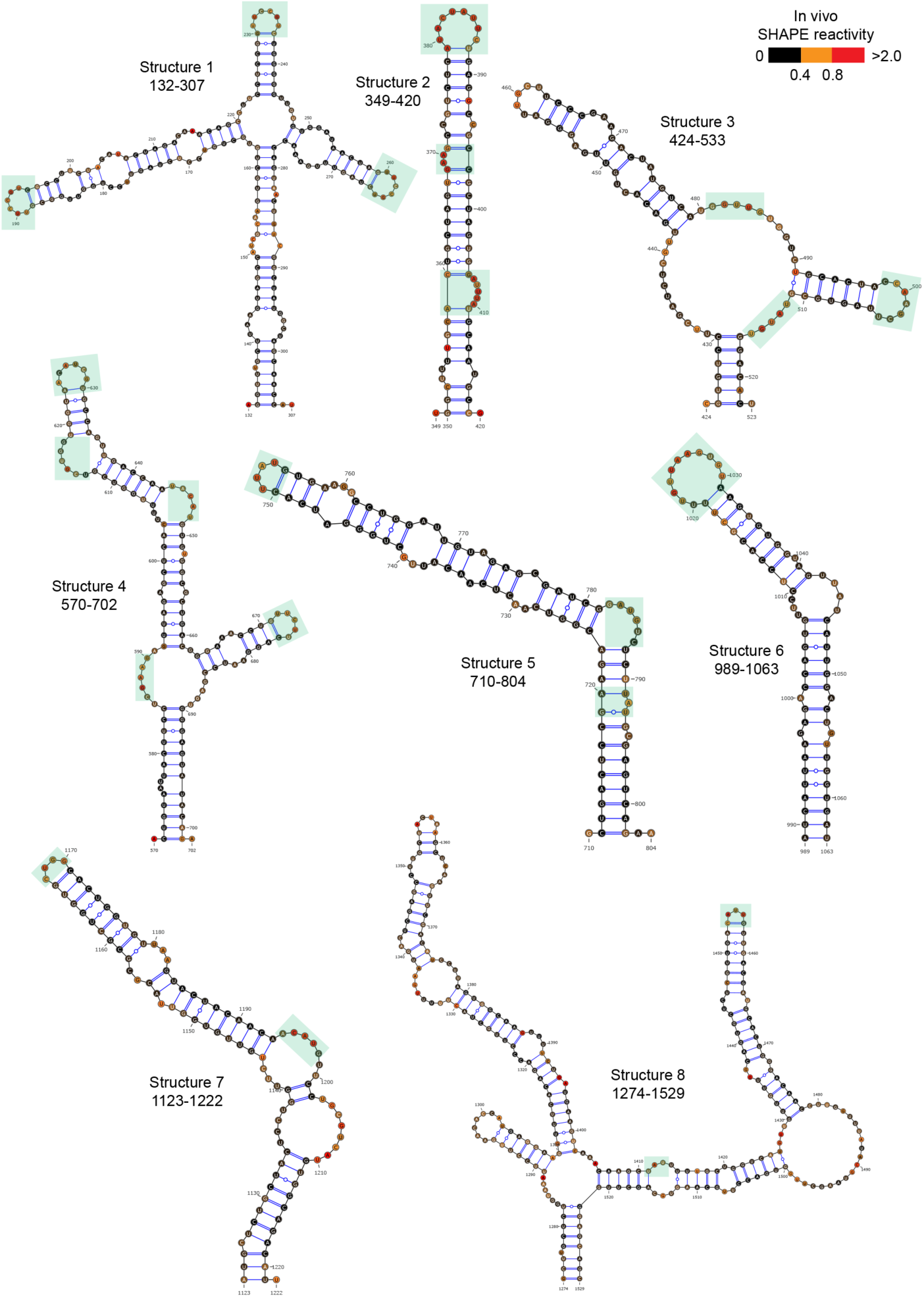
60 well-folded structures (1-8) in PEDV genome. Secondary structure predicted by SHAPE reactivity as a constraint. Green shading indicates high accessibility windows.

**Figure 2—figure supplement 3.**
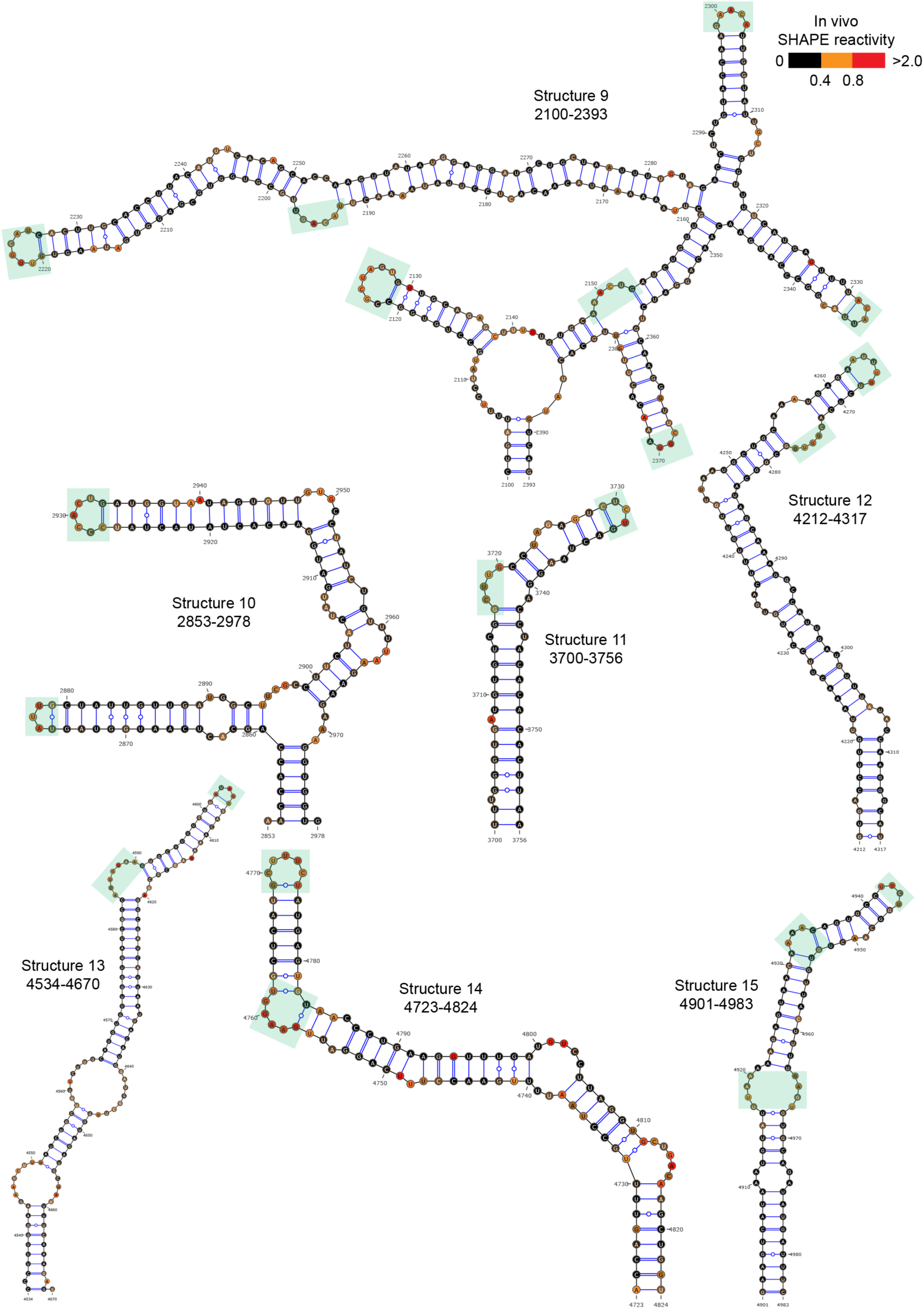
60 well-folded structures (9-15) in PEDV genome. Secondary structure predicted by SHAPE reactivity as a constraint. Green shading indicates high accessibility windows.

**Figure 2—figure supplement 4.**
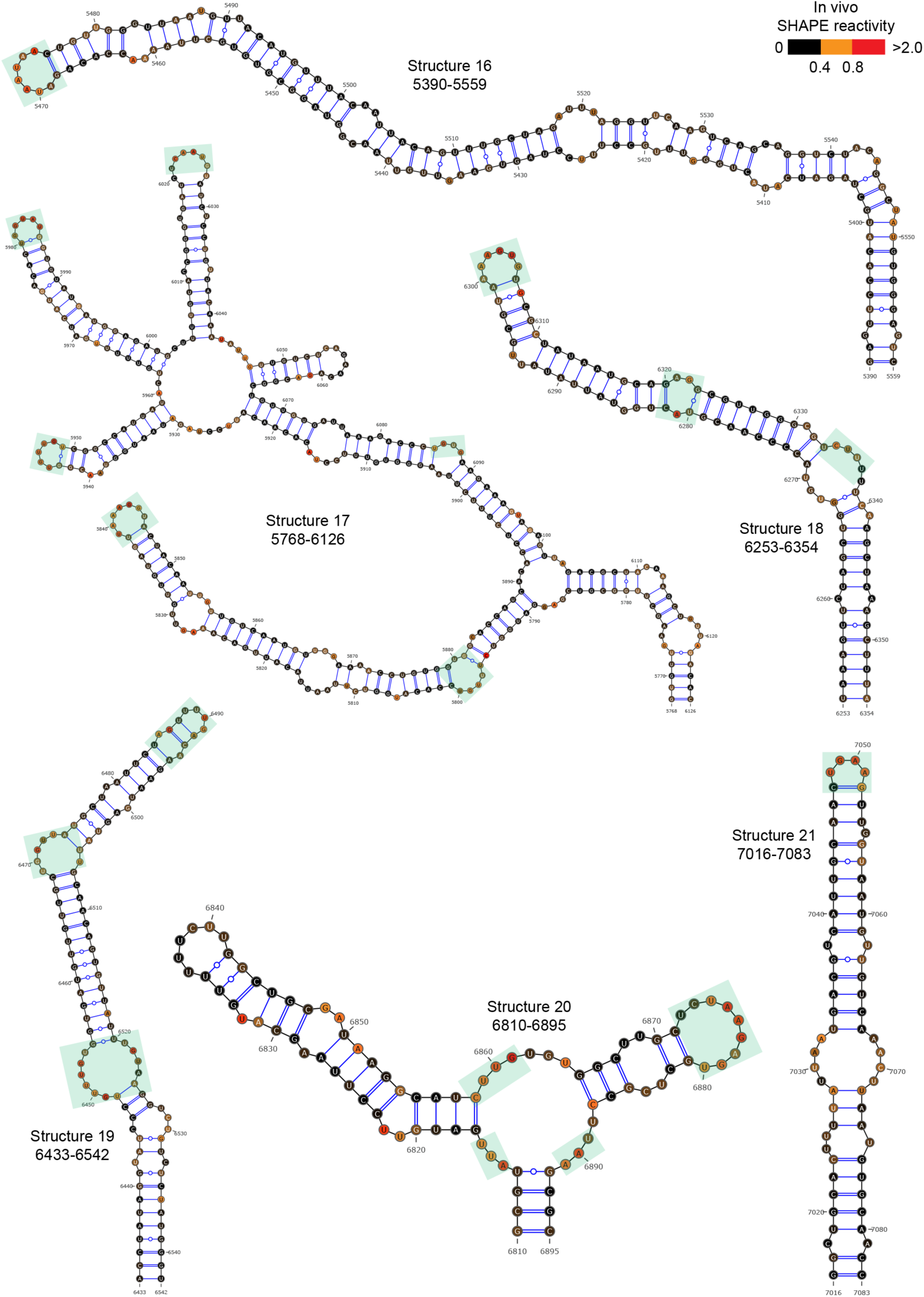
60 well-folded structures (10-21) in PEDV genome. Secondary structure predicted by SHAPE reactivity as a constraint. Green shading indicates high accessibility windows.

**Figure 2—figure supplement 5.**
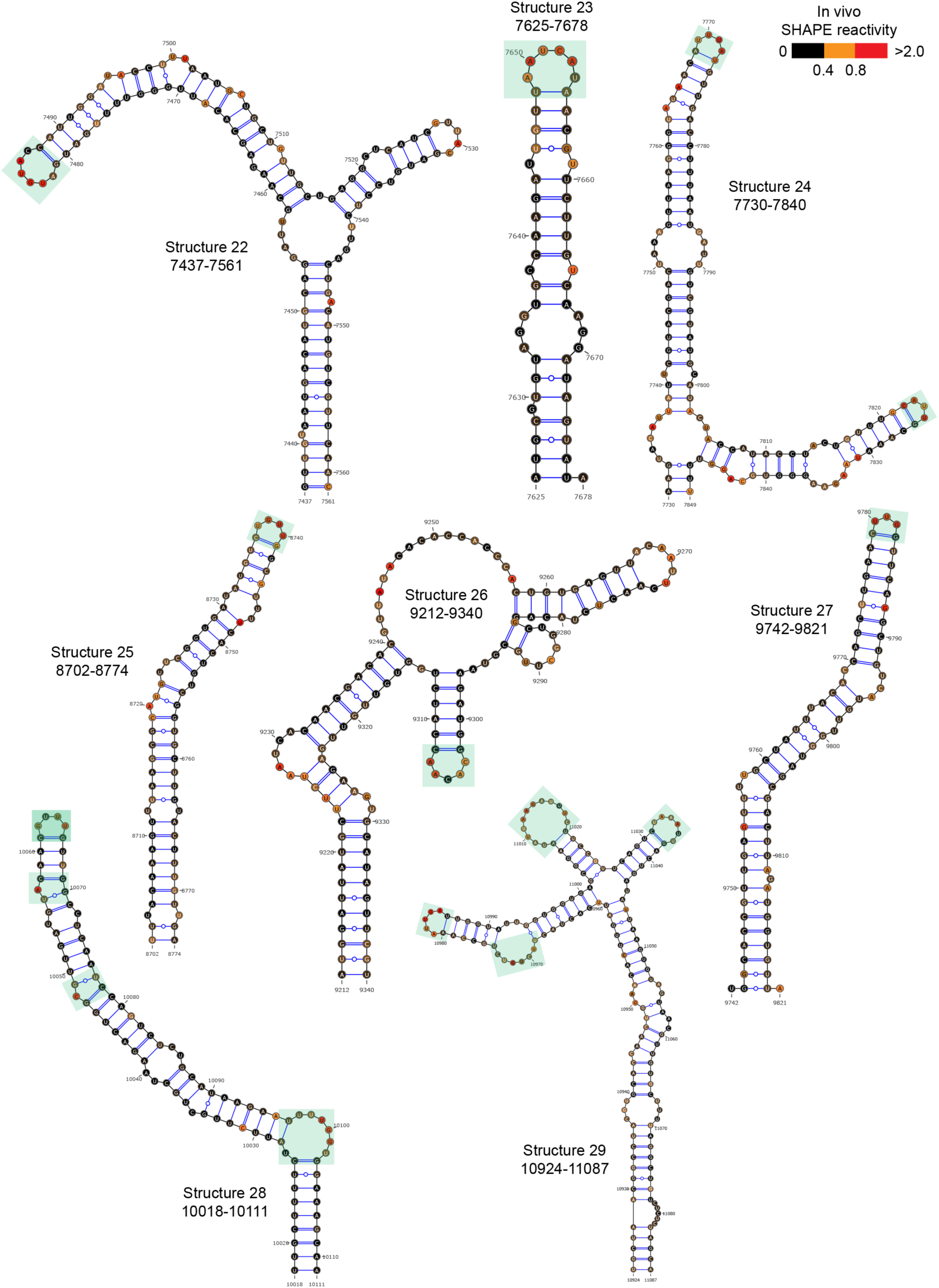
60 well-folded structures (22-29) in PEDV genome. Secondary structure predicted by SHAPE reactivity as a constraint. Green shading indicates high accessibility windows.

**Figure 2—figure supplement 6.**
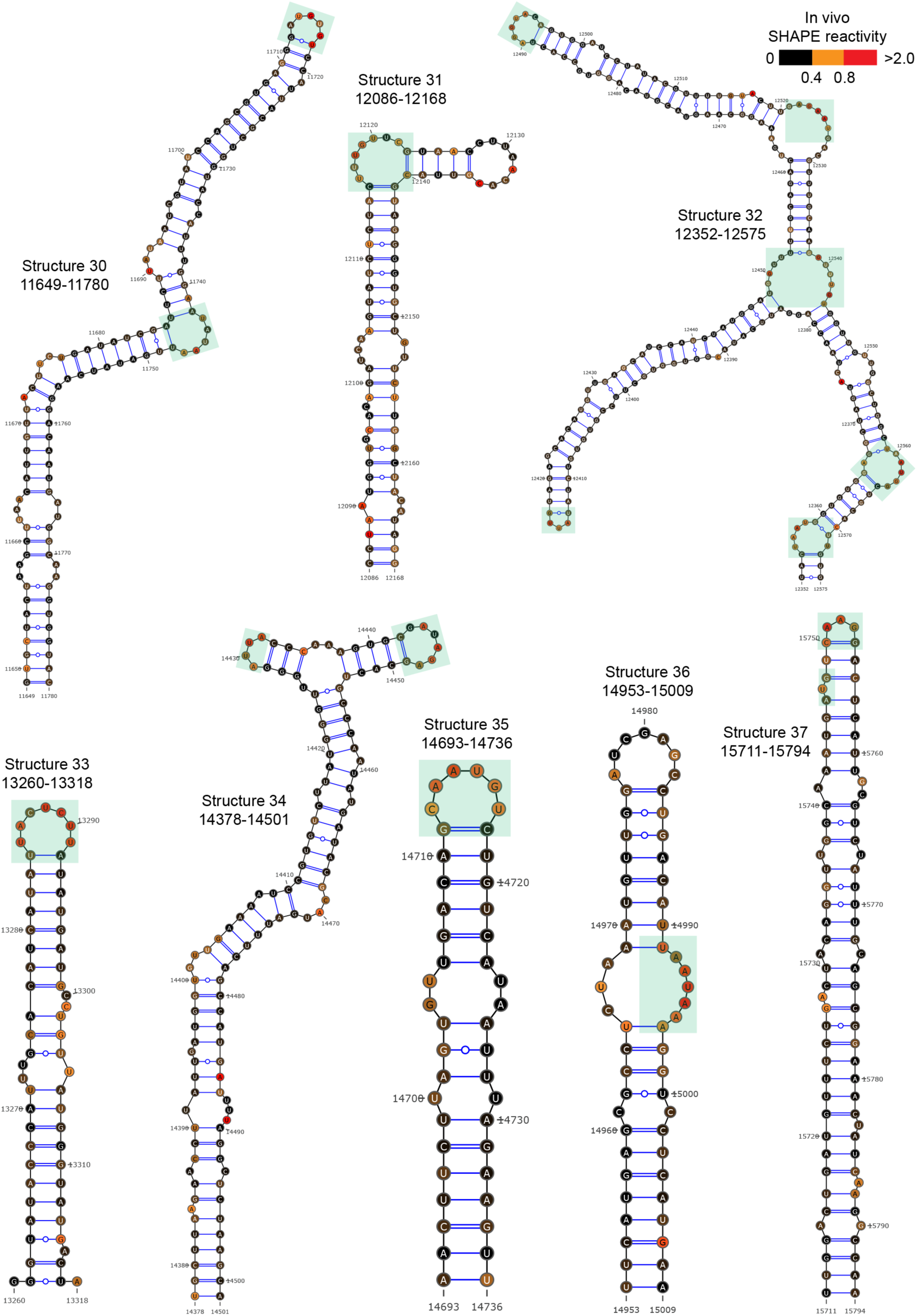
60 well-folded structures (30-37) in PEDV genome. Secondary structure predicted by SHAPE reactivity as a constraint. Green shading indicates high accessibility windows.

**Figure 2—figure supplement 7.**
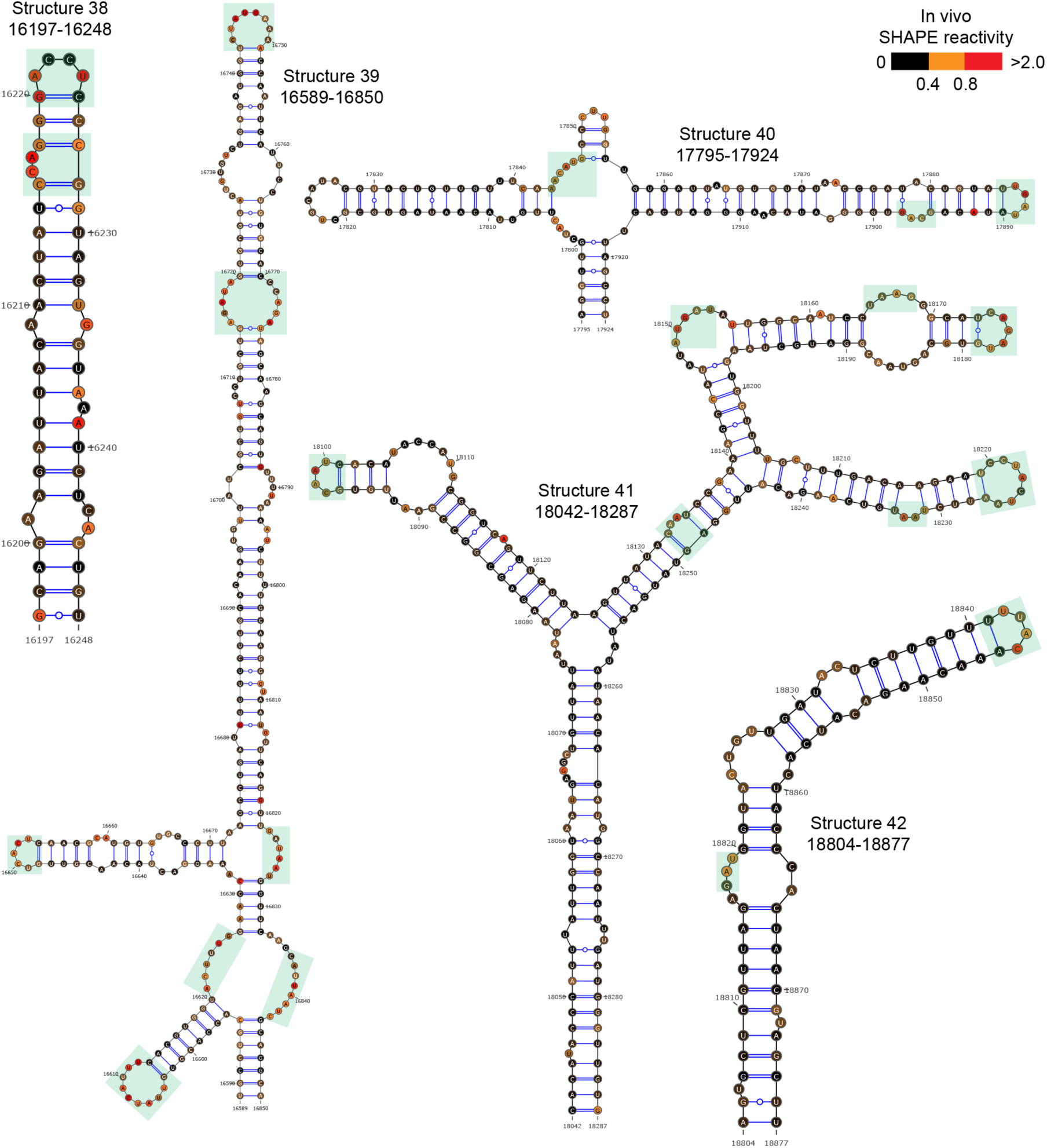
60 well-folded structures (38-42) in PEDV genome. Secondary structure predicted by SHAPE reactivity as a constraint. Green shading indicates high accessibility windows.

**Figure 2—figure supplement 8.**
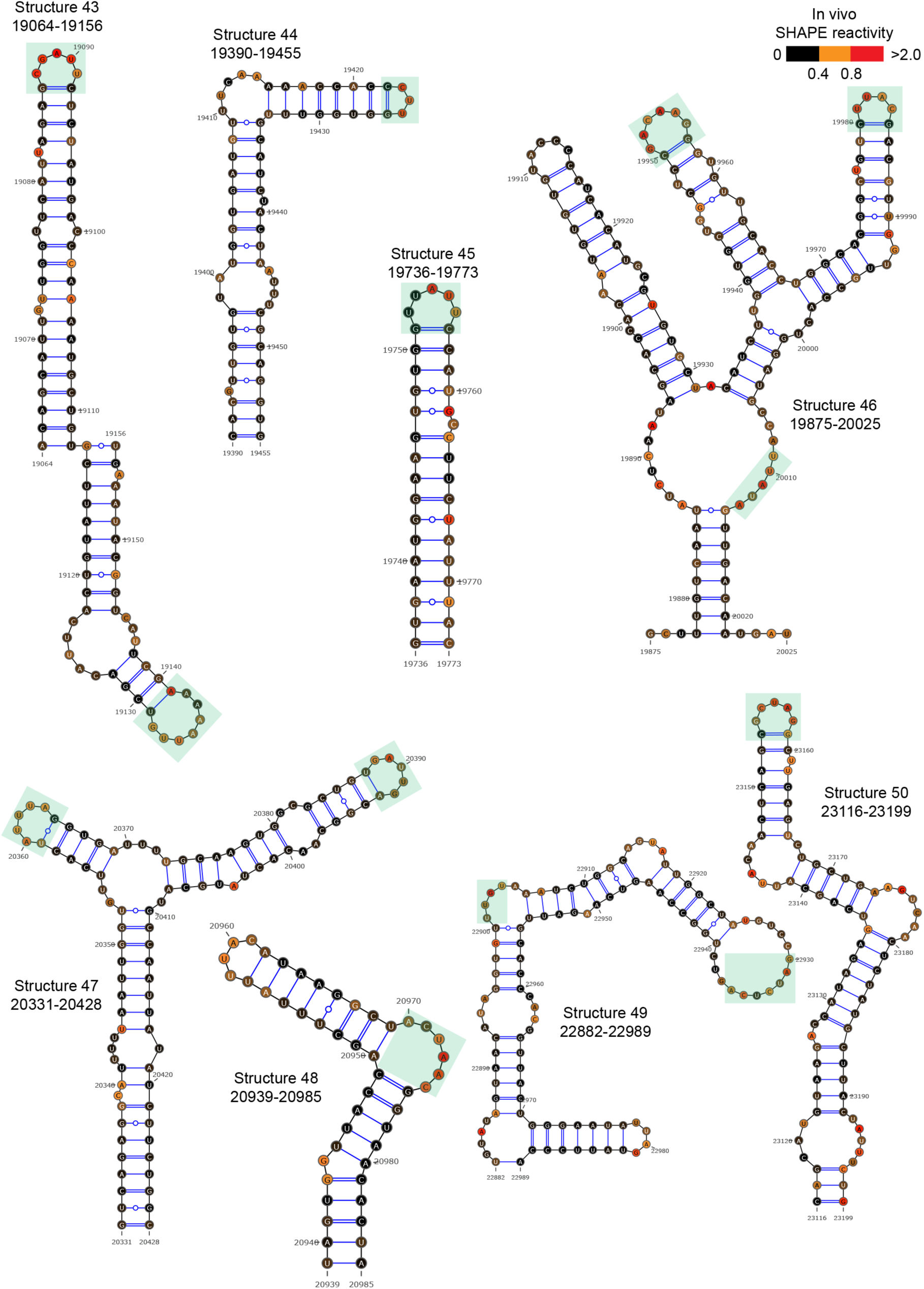
60 well-folded structures (43-50) in PEDV genome. Secondary structure predicted by SHAPE reactivity as a constraint. Green shading indicates high accessibility windows.

**Figure 2—figure supplement 9.**
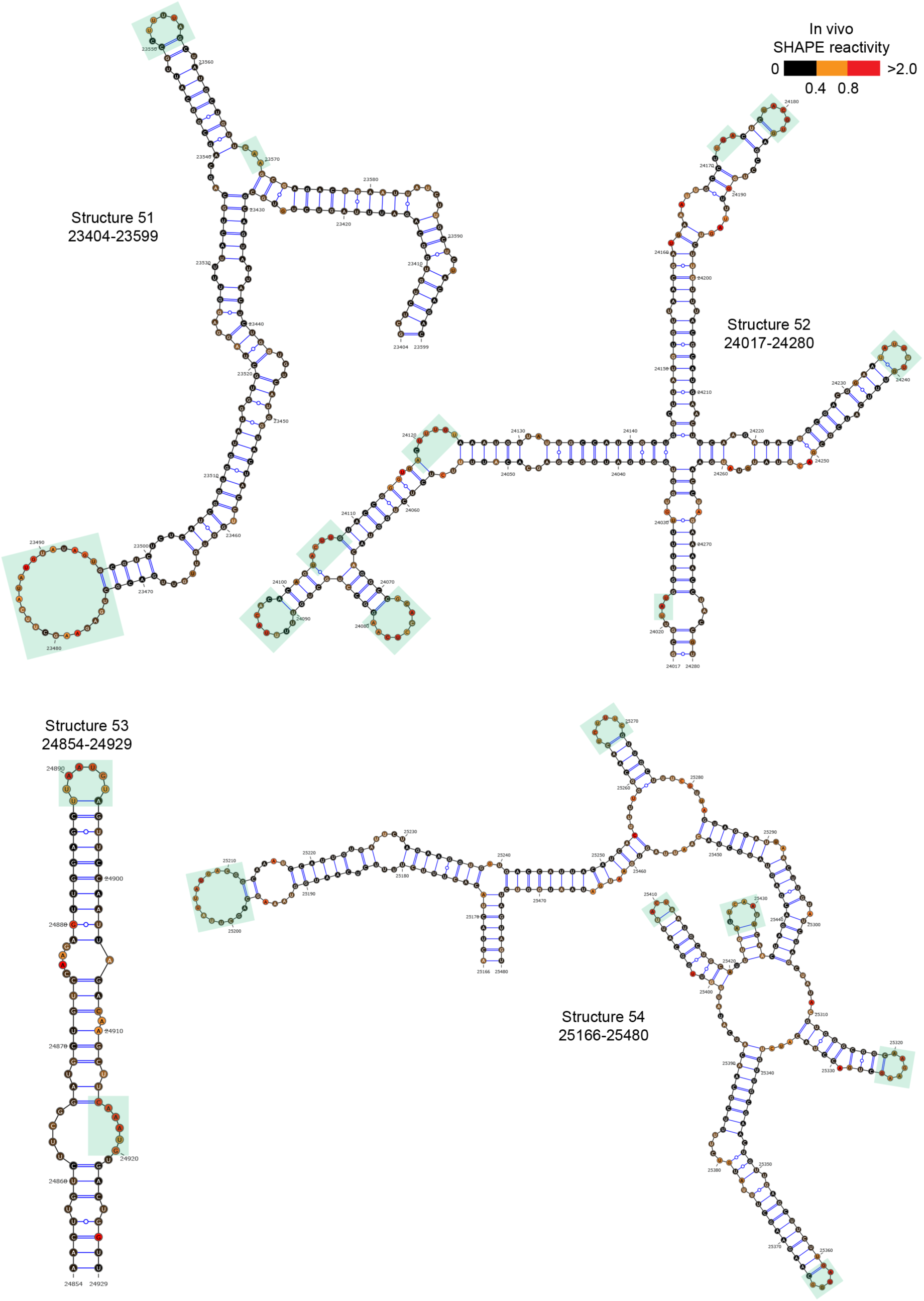
60 well-folded structures (51-54) in PEDV genome. Secondary structure predicted by SHAPE reactivity as a constraint. Green shading indicates high accessibility windows.

**Figure 2—figure supplement 10.**
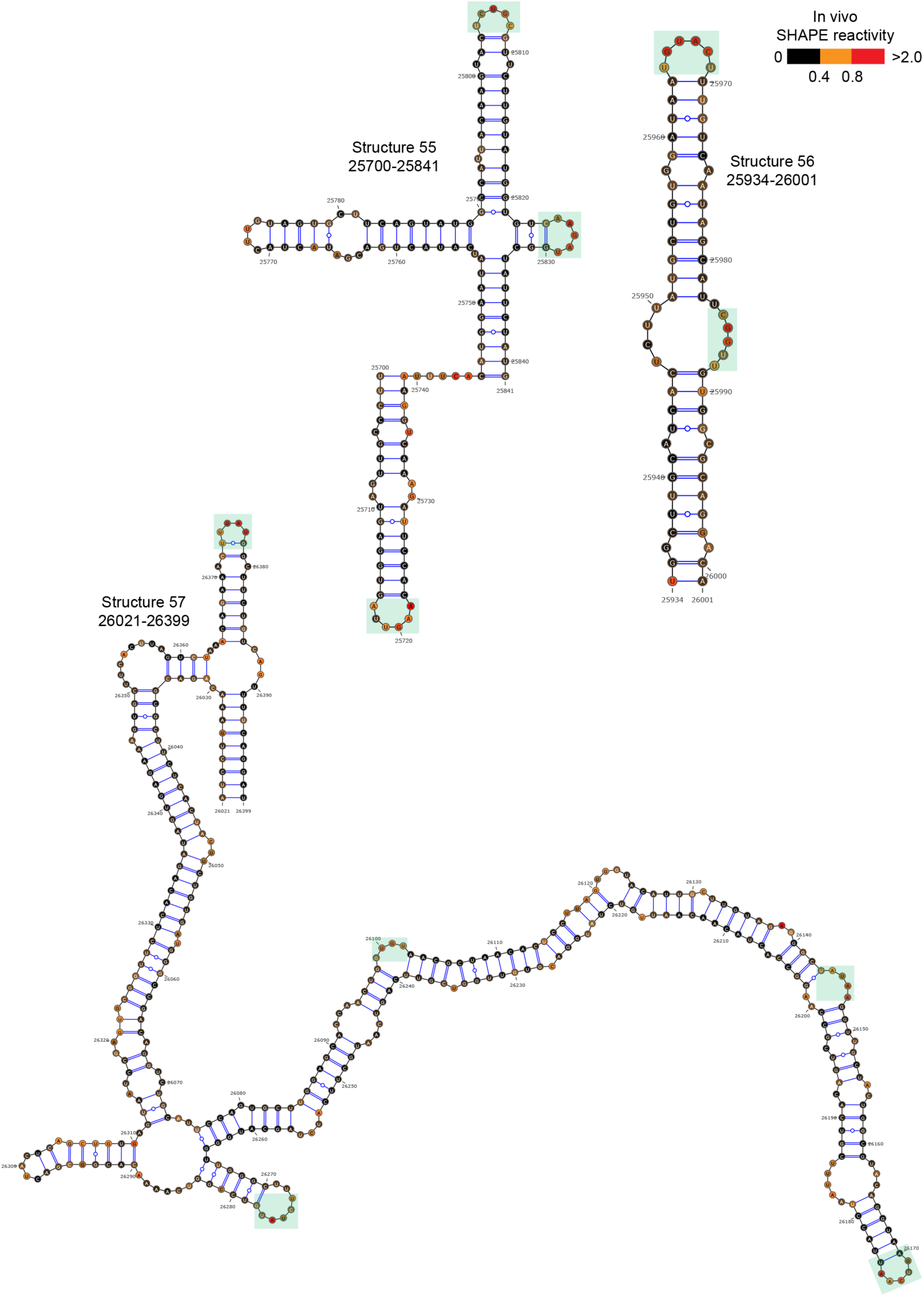
60 well-folded structures (55-57) in PEDV genome. Secondary structure predicted by SHAPE reactivity as a constraint. Green shading indicates high accessibility windows.

**Figure 2—figure supplement 11.**
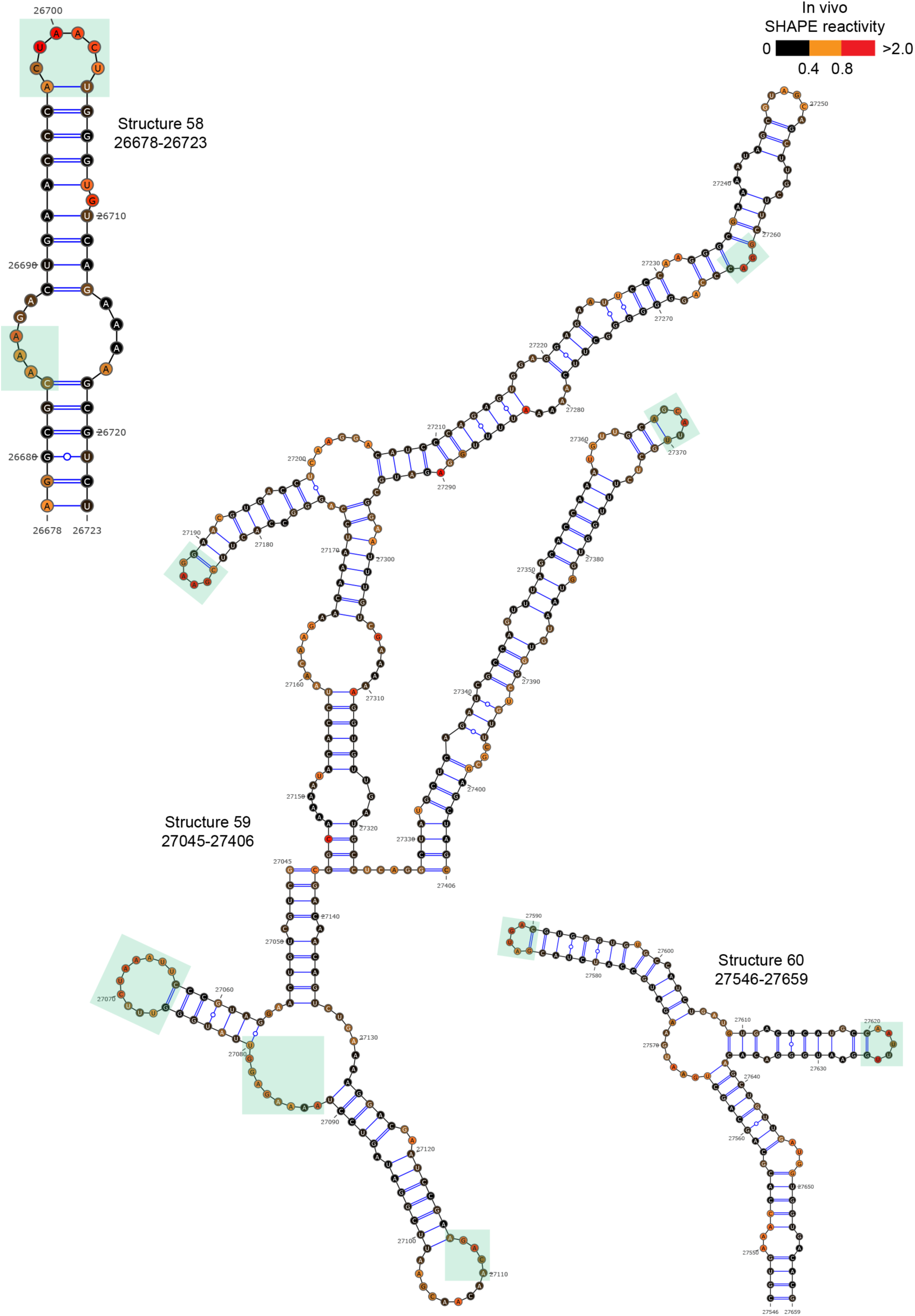
60 well-folded structures (58-60) in PEDV genome. Secondary structure predicted by SHAPE reactivity as a constraint. Green shading indicates high accessibility windows.

**Figure 2—figure supplement 12.**
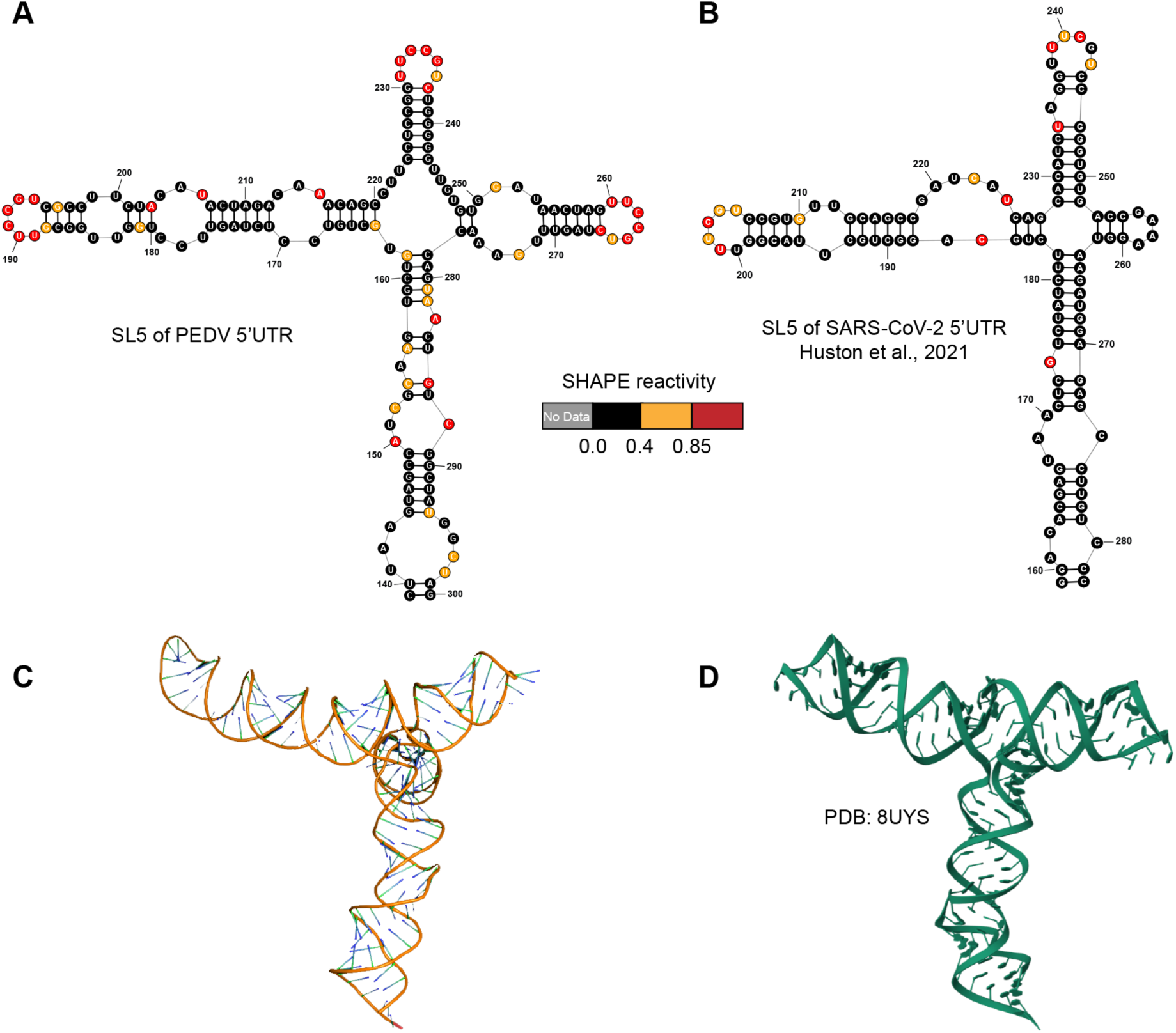
Structures of 5’UTR-SL5 in the genomes of PEDV and SARS-CoV-2. **(A)** Secondary structure of 5’UTR-SL5 in the PEDV genome predicted by SHAPE reactivity as a constraint. **(B)** Secondary structure of 5’UTR-SL5 in the SARS-CoV-2 genome (Huston et al., 2021). **(C)** The three-dimensional structural model of PEDV 5’UTR-SL5 based on secondary structure was predicted using FARFAR2 (Watkins et al., 2020). **(D)** Three-dimensional of 5’UTR-SL5 in the SARS-CoV-2 genome (PDB id: 8UYS).

**Figure 2—figure supplement 13.**
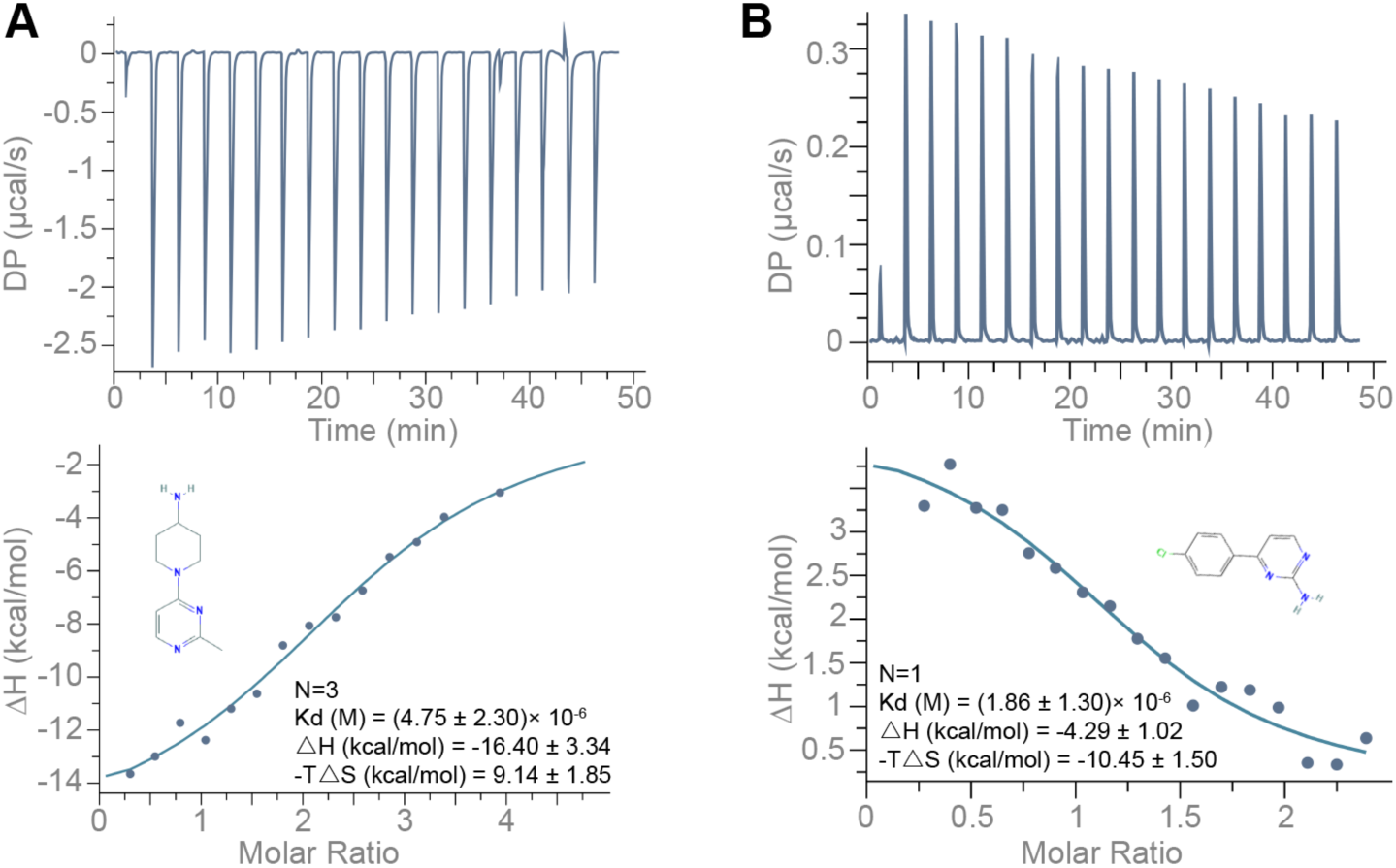
ITC Binding Profile of **(A)** Compound 1 (1-(2-methylpyrimidin-4-yl) piperidin-4-amine; 250 μM) and **(B)** Compound 4 (4-(4-Chlorophenyl) pyrimidin-2-amine; 250 μM) titrated into the 10 μM 5’UTR-SL5 of PEDV. Buffer solution: 100 mM HEPES (pH 8.0), 100 mM NaCl and 10 mM MgCl2. Temperature: 25℃.

**Figure 2—figure supplement 14.**
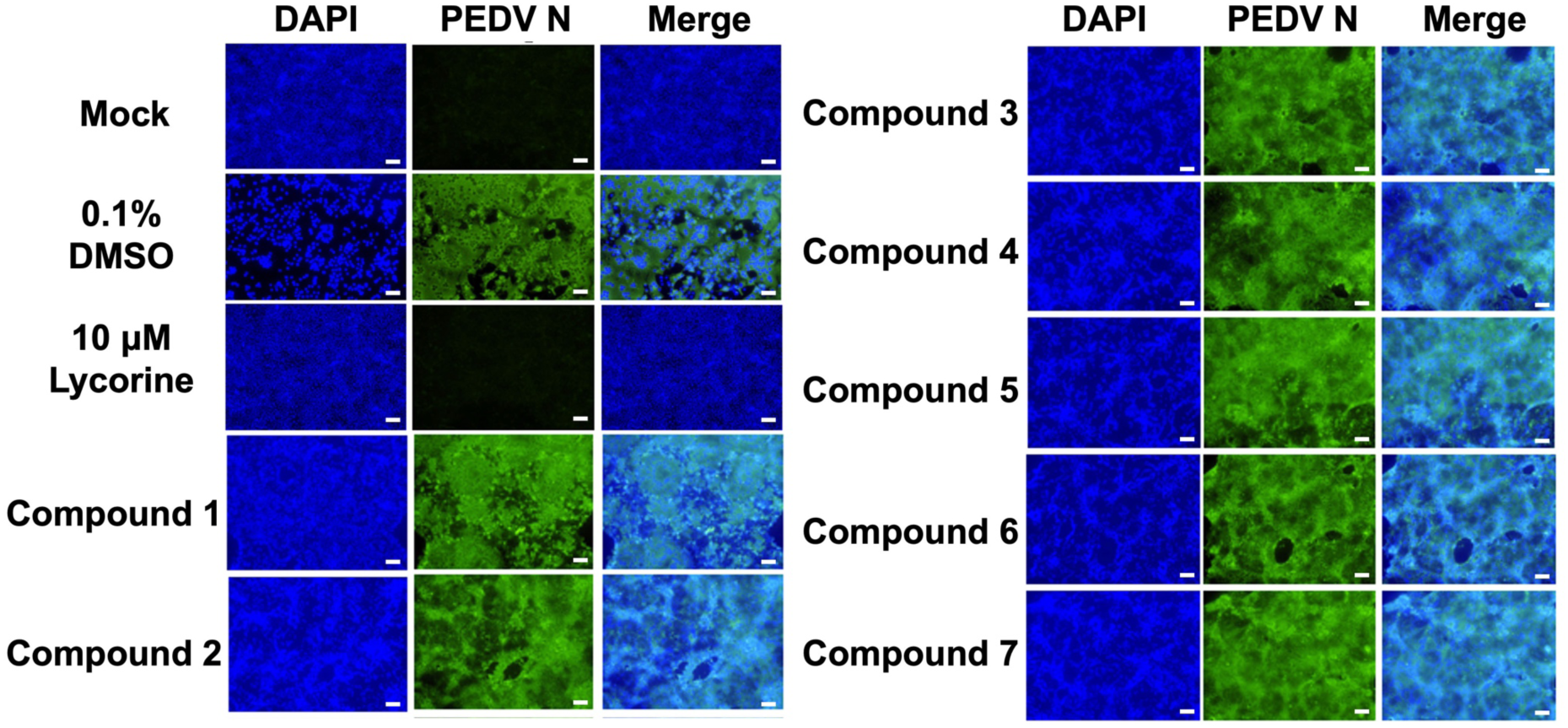
Antiviral activity of compounds interacting with the PEDV 5’UTR-SL5 *in vitro*. Lycorine is a PEDV-positive inhibitor. Confocal fluorescence microphotographs of the same cells with DAPI (left), N protein of PEDV (middle) or Merge (right) were demonstrated (Scale bar = 50 μm).

**Figure 3—figure supplement 1.**
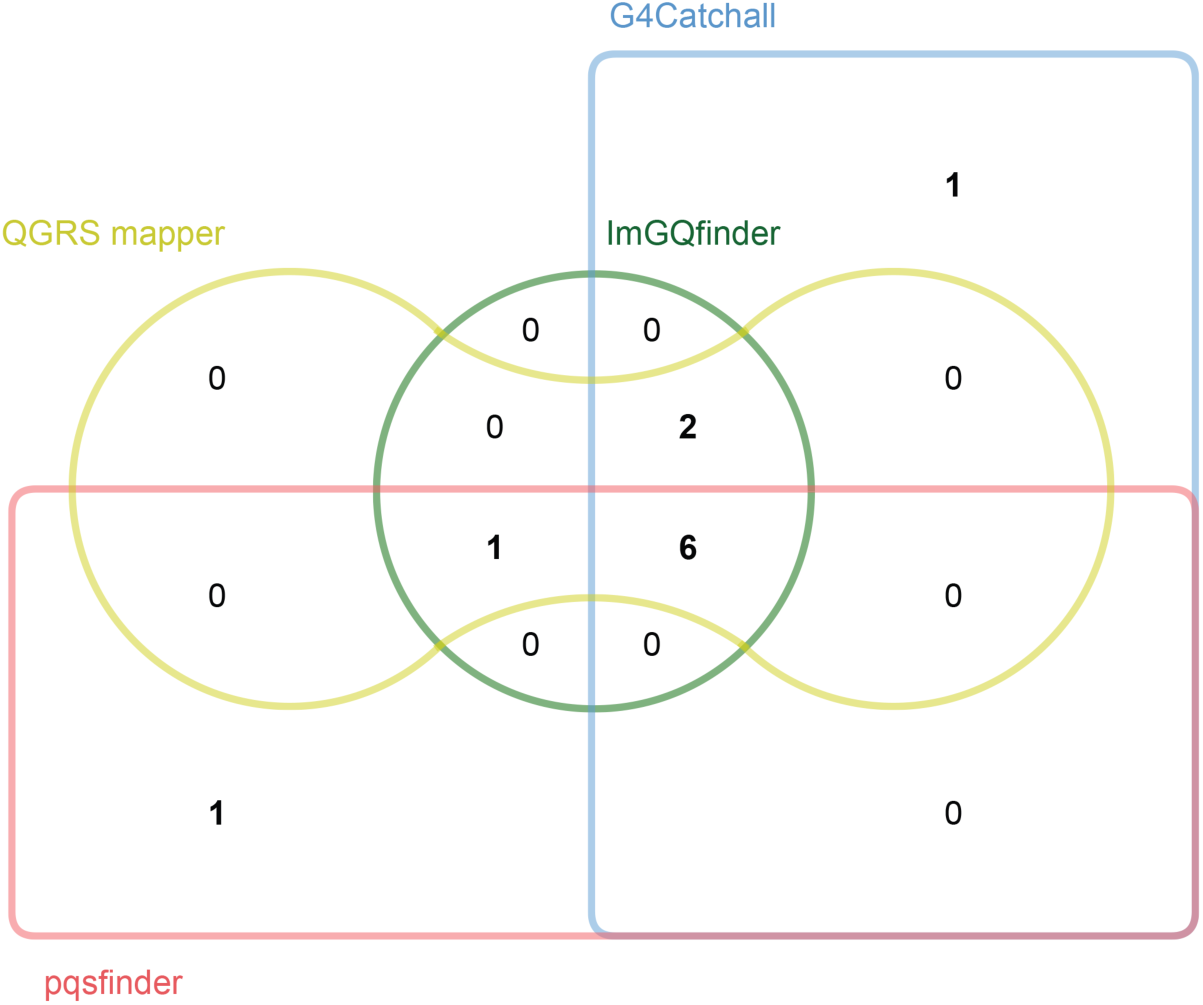
Prediction of potential G-quadruplex forming sequences (PQSs) in the PEDV genome by different prediction tools. Edwards’ Venn diagram showing the overlap of the putative potential G-quadruplex sequences (PQSs) predicted by G4Catchall, QGRS mapper, ImGQfinder and pqsfinder.

**Figure 3—figure supplement 2.**
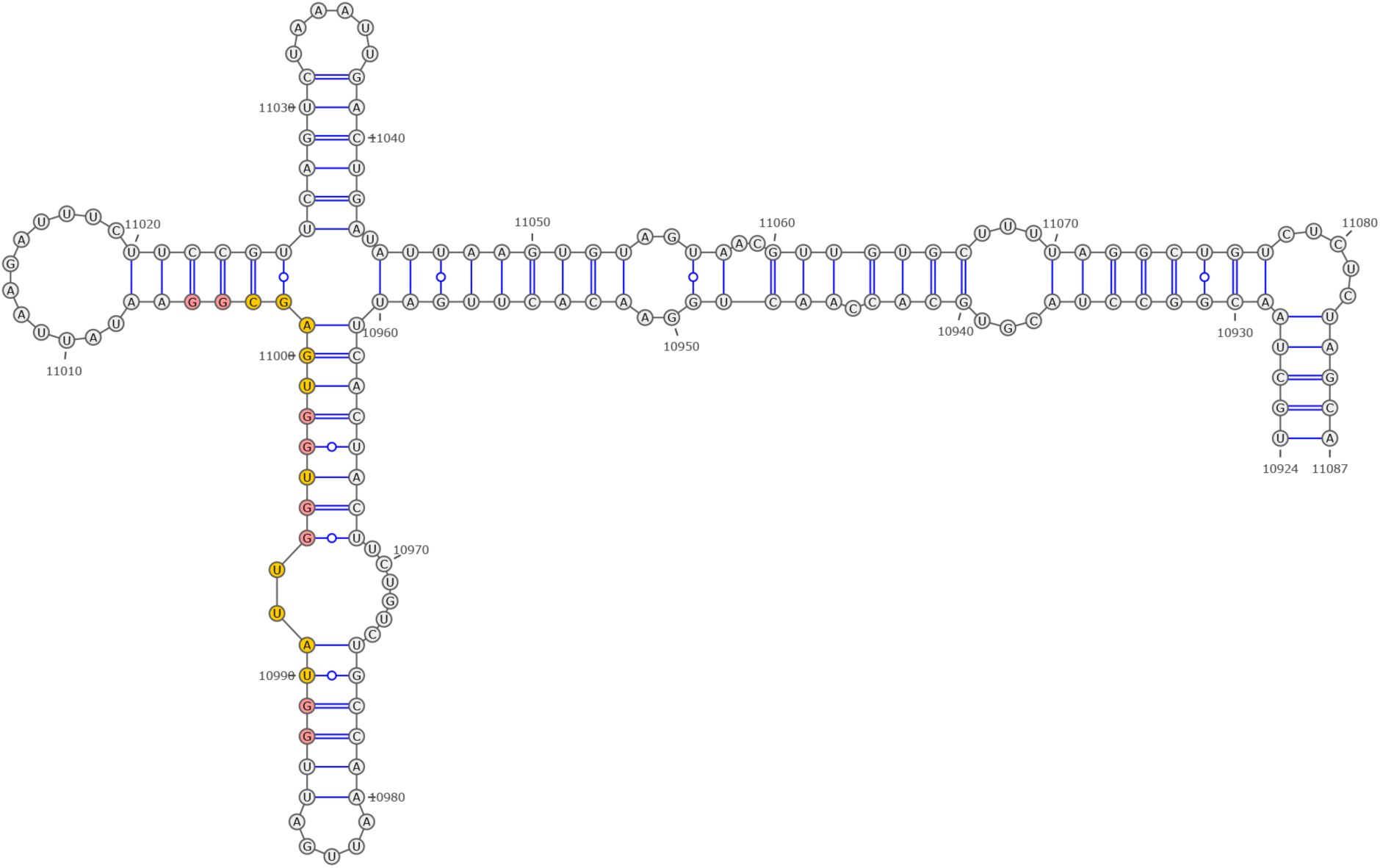
Secondary structure of the local region containing PQS2 predicted by SHAPE reactivity as a constraint.

**Figure 3—figure supplement 3.**
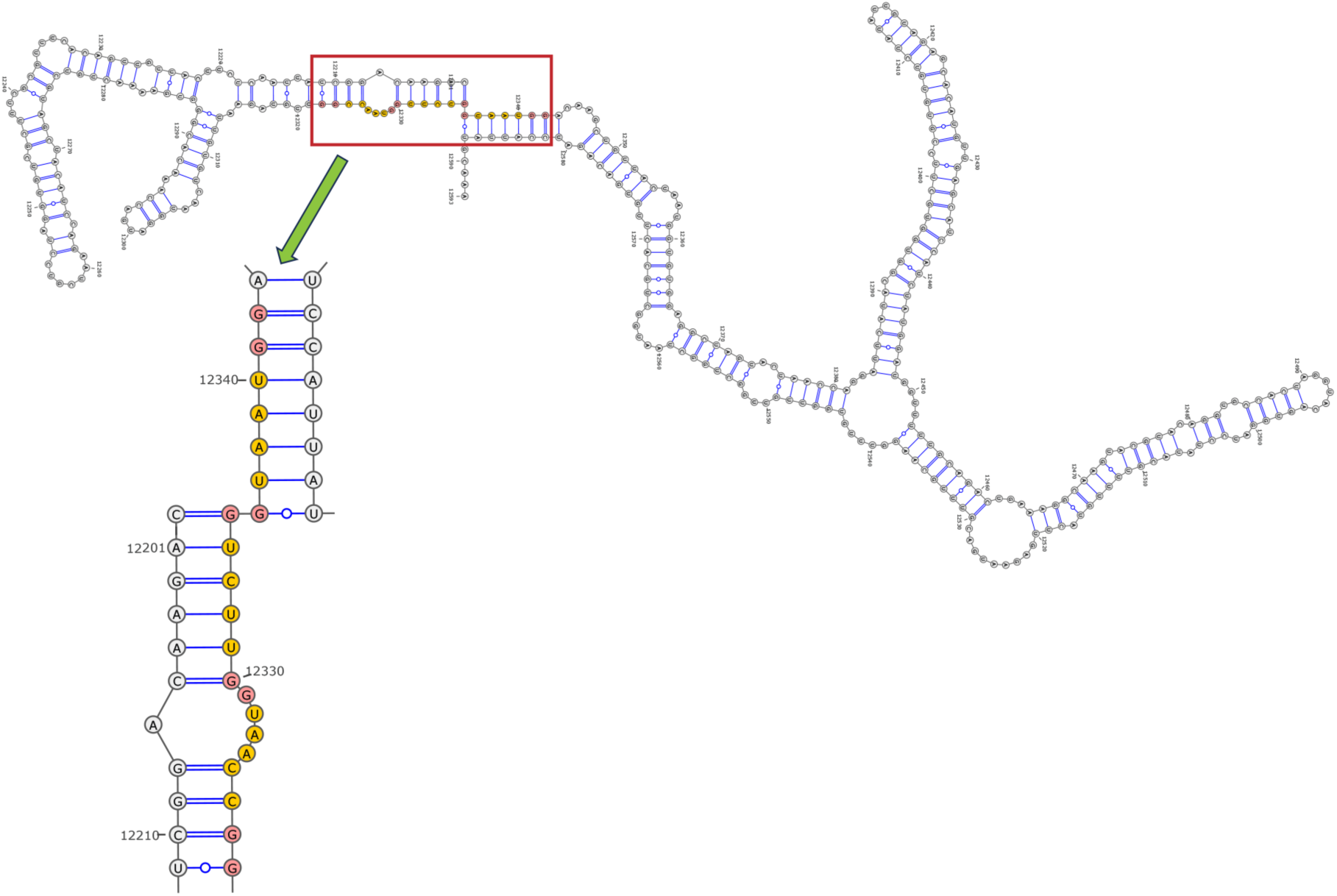
Secondary structure of the local region containing PQS3 predicted by SHAPE reactivity as a constraint.

**Figure 3—figure supplement 4.**
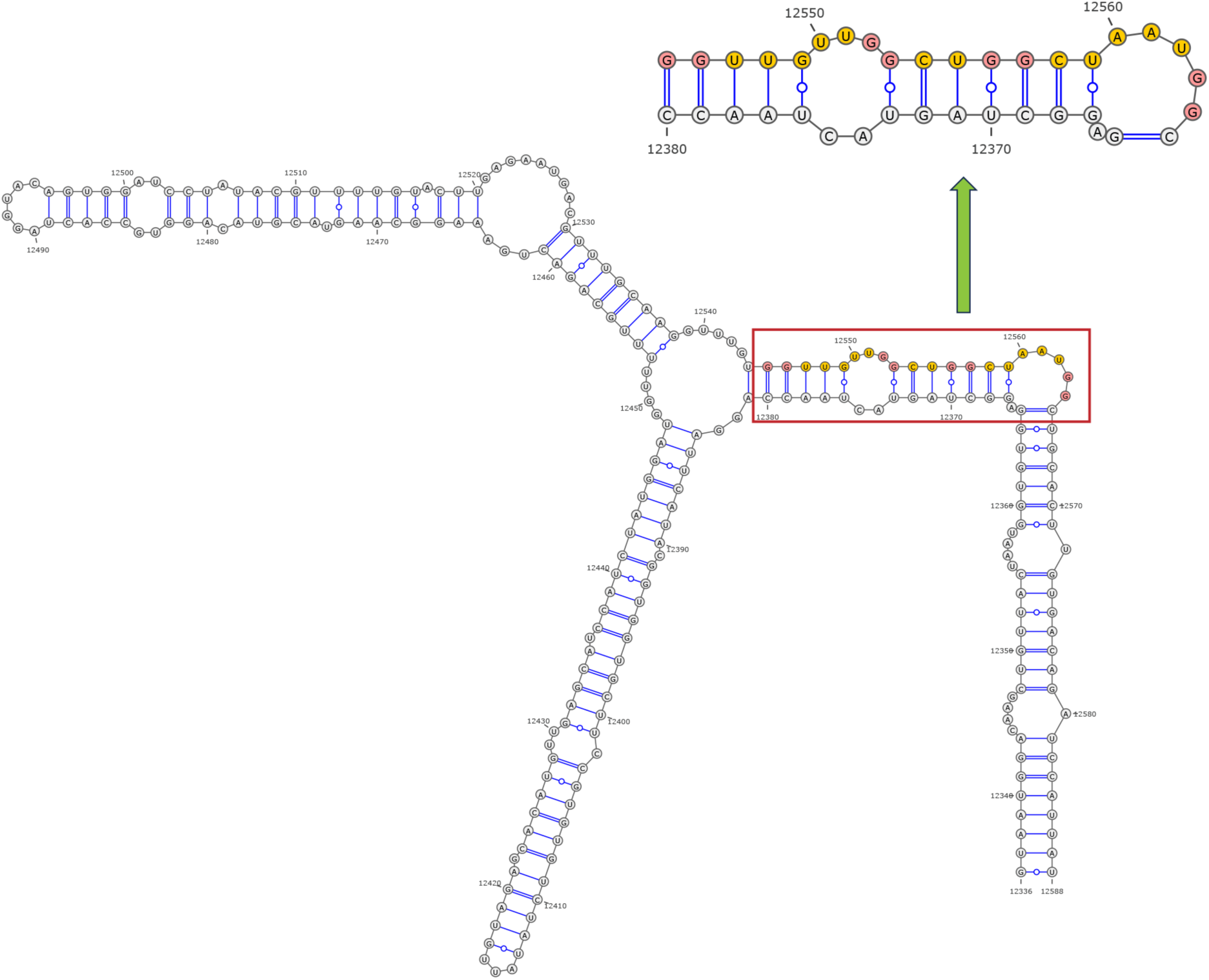
Secondary structure of the local region containing PQS4 predicted by SHAPE reactivity as a constraint.

**Figure 3—figure supplement 5.**
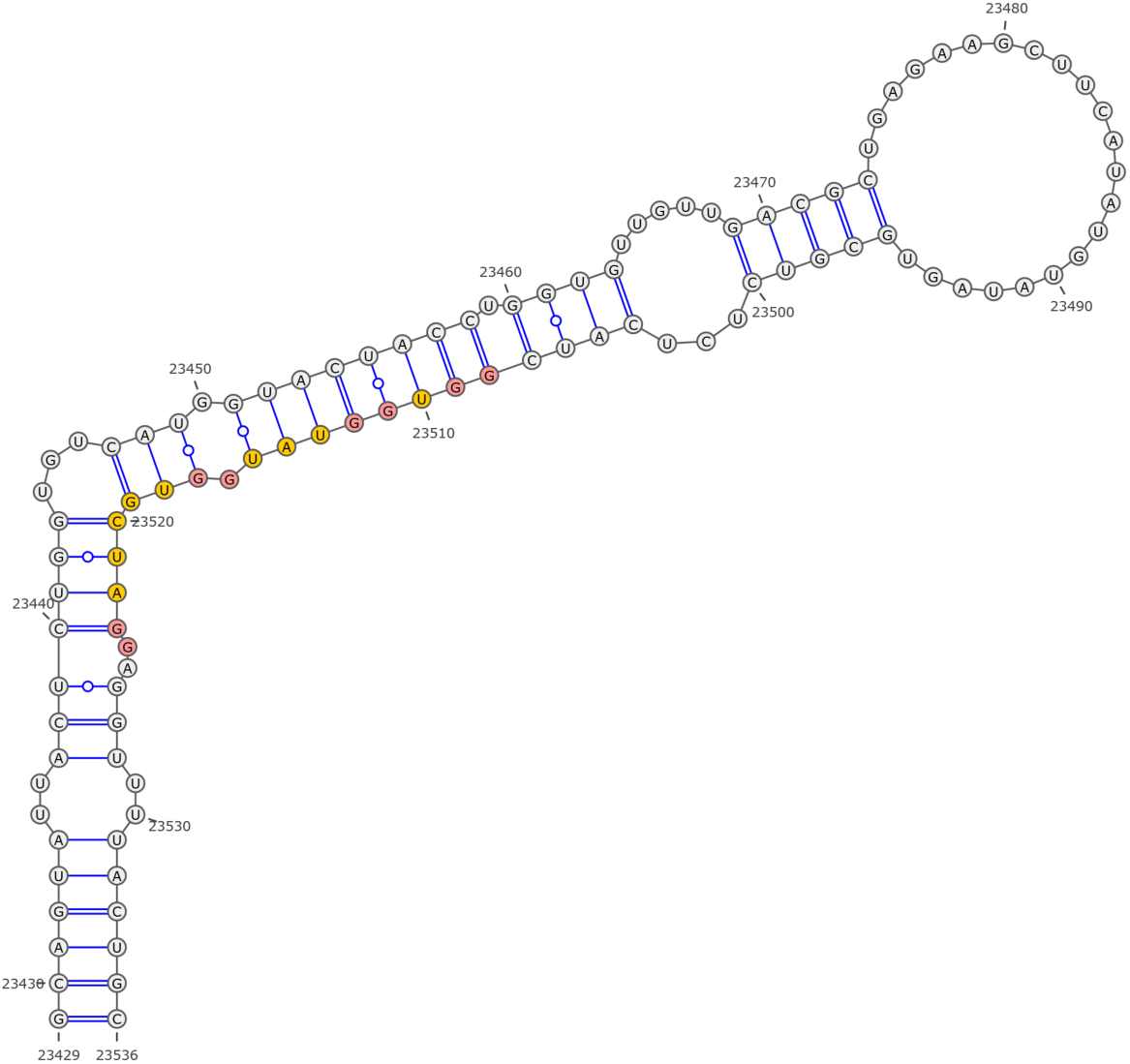
Secondary structure of the local region containing PQS6 predicted by SHAPE reactivity as a constraint.

**Figure 3—figure supplement 6.**
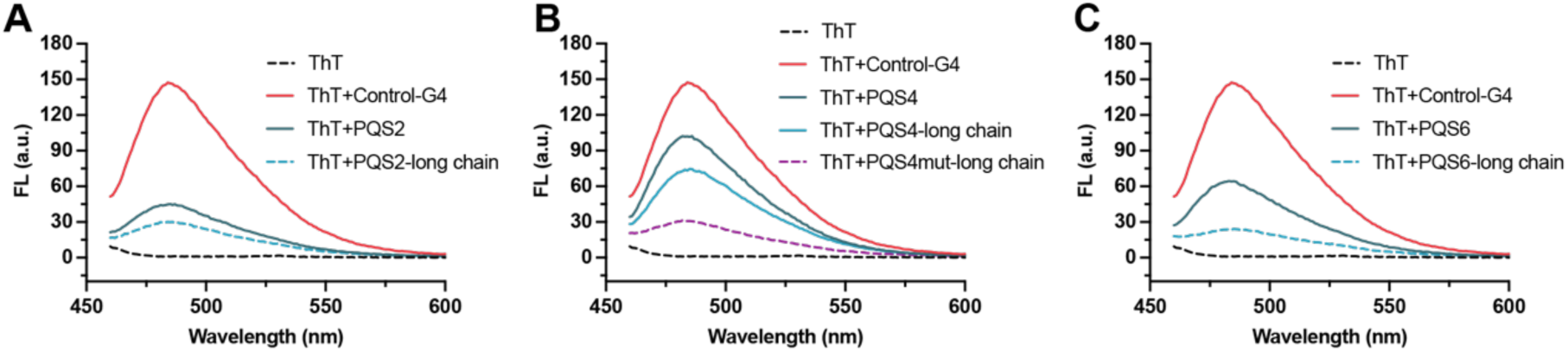
**(A)** Fluorescence turn-on assays of ThT (1 μM) in the presence of PQS2 (0.5 μM) and PQS2-long chain (0.5 μM). **(B)** Fluorescence turn-on assays of ThT (1 μM) in the presence of PQS4 (0.5 μM), PQS4-long chain (0.5 μM) and PQS4mut-long chain (0.5 μM). **(C)** Fluorescence turn-on assays of ThT (1 μM) in the presence of PQS6 (0.5 μM) and PQS6-long chain (0.5 μM). Excitation wavelength was 442 nm. PRRSV-G4 RNA was used as control.

**Figure 3—figure supplement 7.**
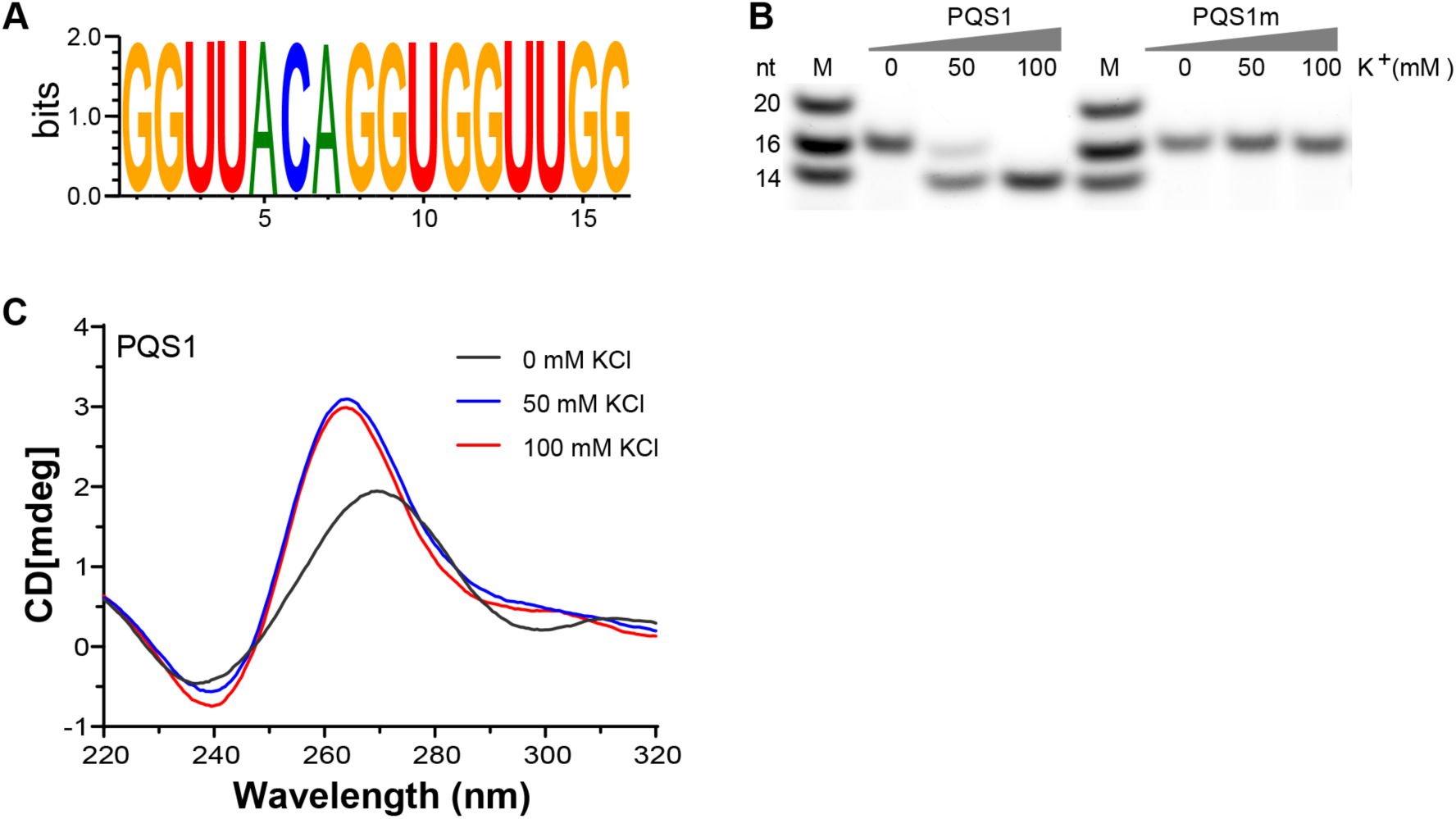
G-quadruplex structure folded by PQS1. **(A)** WebLogo representation nucleotide conservation of PQS1 among different PEDV strains. The height of the letters indicates the relative frequency of occurrence of a particular nucleotide at that position. **(B)** Native-PAGE analysis of the migration of G4 RNAs (PQS1) and G4 mutant RNAs (PQS1m) was performed under different KCl concentrations. Lanes 1 and 5, RNA ladder; lanes 2, 3, 4, PQS1; lanes 6, 7, 8, PQS1m. **(C)** CD spectroscopy of PQS1 in the presence of different concentrations of KCl.

**Figure 4—figure supplement 1.**
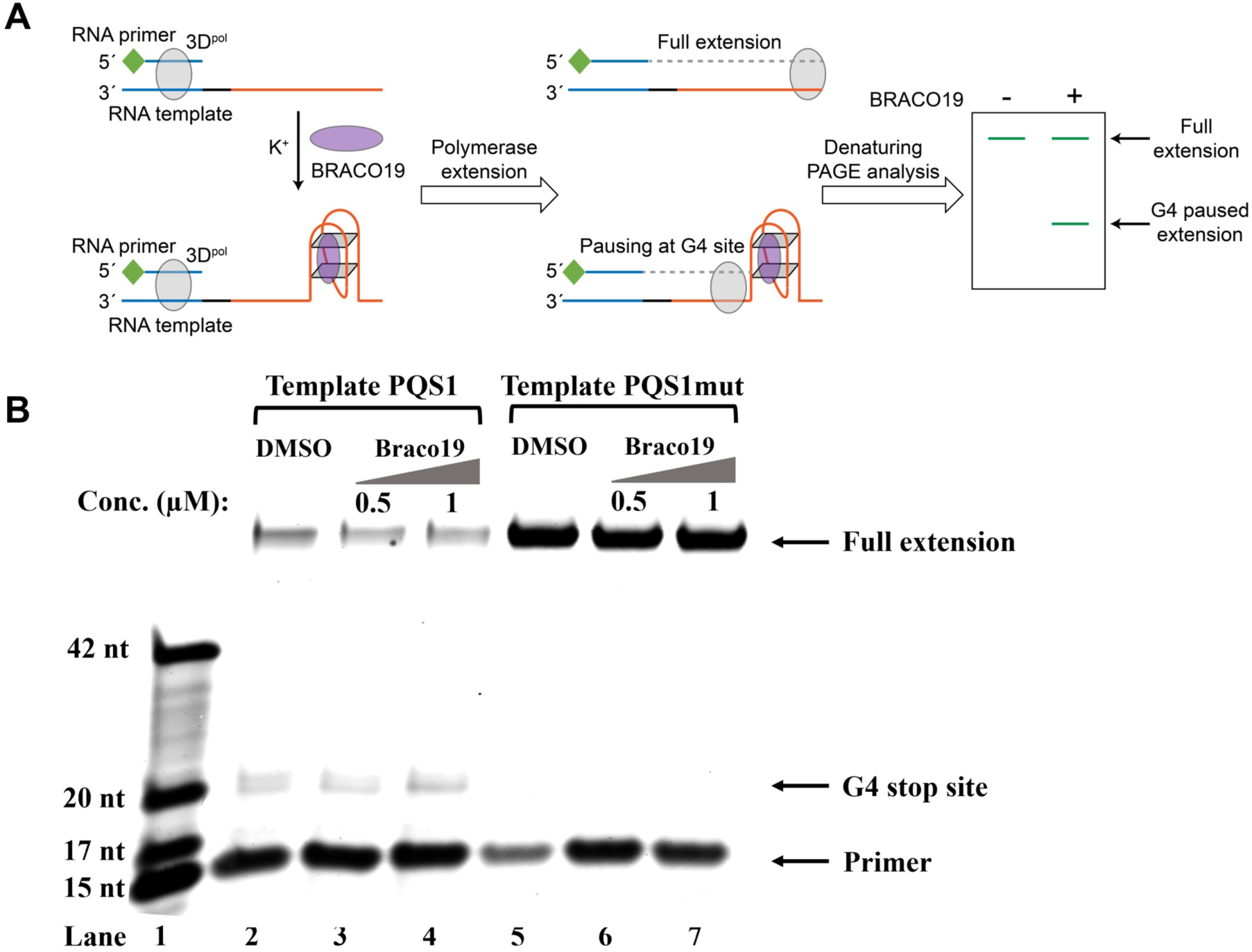
G4 structure of PQS1 inhibits RNA-dependent RNA synthesis. **(A)** Schematic diagram of the RNA stop assay. The PQS1 sequence (orange part) was integrated into the 5’ end of the RNA template, and a 5’ FAM-tagged RNA primer P15 (green and blue parts of the RNA primer) was designed to target the 3’ end of the template (blue part of the RNA blue part of the template) to initiate RNA synthesis. When G4 blocks RNA synthesis, there are fewer full-length elongated bands and truncated bands at the site of G4 formation. **(B)** Denaturing-PAGE analysis. Black arrows indicate the positions of the full-length product, G4 paused product, and free primer. The fully extended product and the template RNA and the polymerase form a stable complex that moves much more slowly than the ssRNA. Partial full-length extended products and stopped products (due to G-quadruplex fold) were observed along template PQS1 (lane 2), but only fully extended products were observed along G4-mutated template PQS1mut (lane 5). When increasing amounts of compound Braco-19 were incubated with template PQS1, a gradual decrease in fully extended products was observed (lanes 2-4). On the contrary, the fully extended products of template PQS1mut were not affected by the addition of Braco-19, and the G4-specific termination event was not characterized (lanes 5-7). Lane 1, RNA ladder (p15, m17, m20 and m42 in Table S1); lanes 2, no Braco19 control; lanes 3 and 4, 0.5 and 1 µM compounds inhibit RNA extension; lanes 5, no Braco19 control; lanes 6 and 7, compounds do not inhibit RNA extension.

**Figure 4—figure supplement 2.**
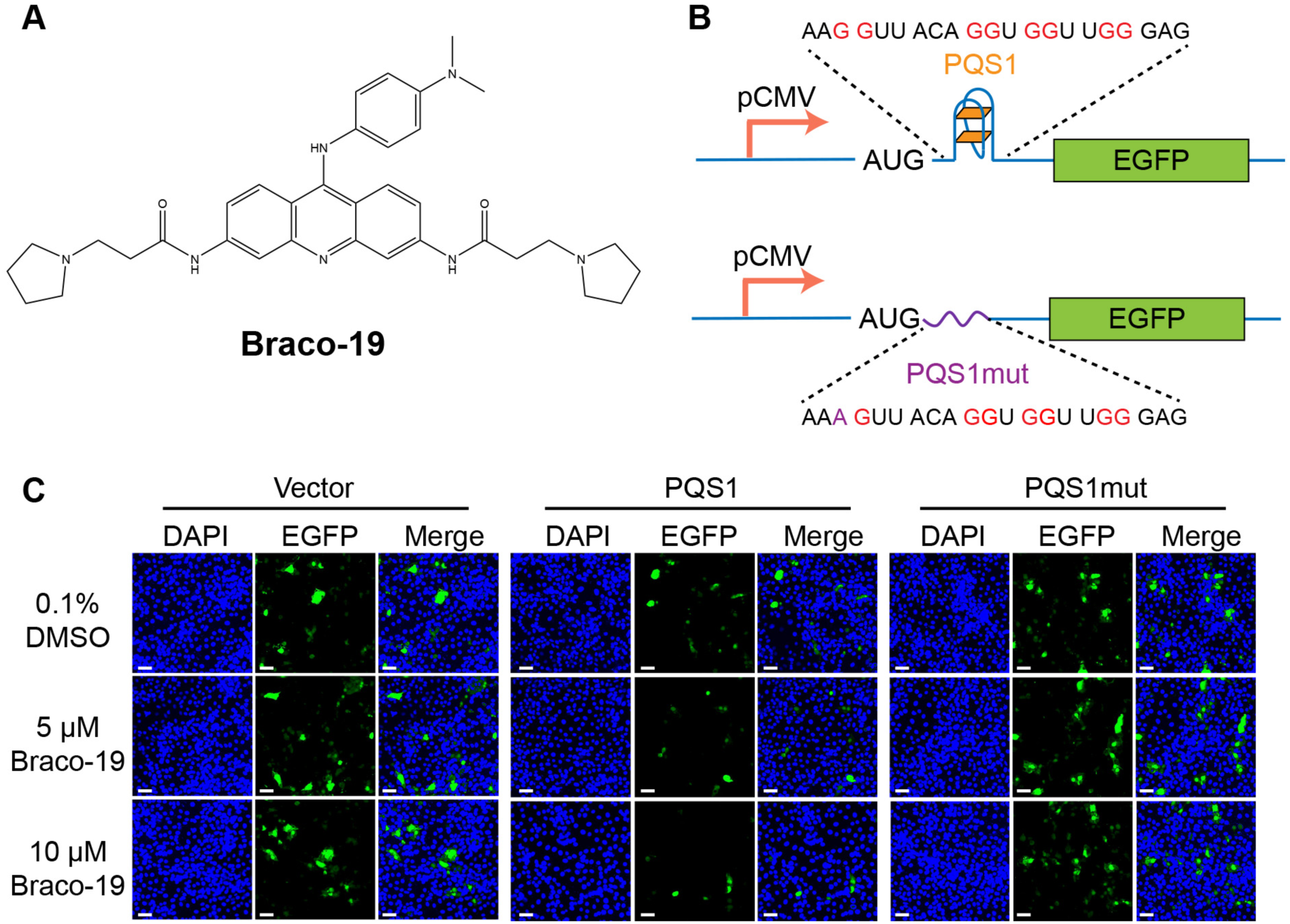
**(A)** Structure of compound Braco-19. **(B)** Schematic representation of the pEGFP-C1 plasmids. The empty vector carries the wild-type EGFP sequence, while the engineered plasmids contain the PEDV PQS1 sequence and the PQS1mut sequence. **(C)** Plasmids containing wild-type EGFP, PQS1 and PQS1mut were transfected into Vero cells, which were then treated with 0, 5 or 10 μM Braco-19 for 24 h. EGFP expression levels were measured via confocal fluorescence microscopy. Compared with DMSO treatment, the addition of Braco-19 led to a significant decrease in PQS1 EGFP fluorescence intensity. In contrast, no difference was observed in the EGFP expression of the PQS1mut vector and the empty vector. Scale bar = 50 μm.

**Figure 4—figure supplement 3.**
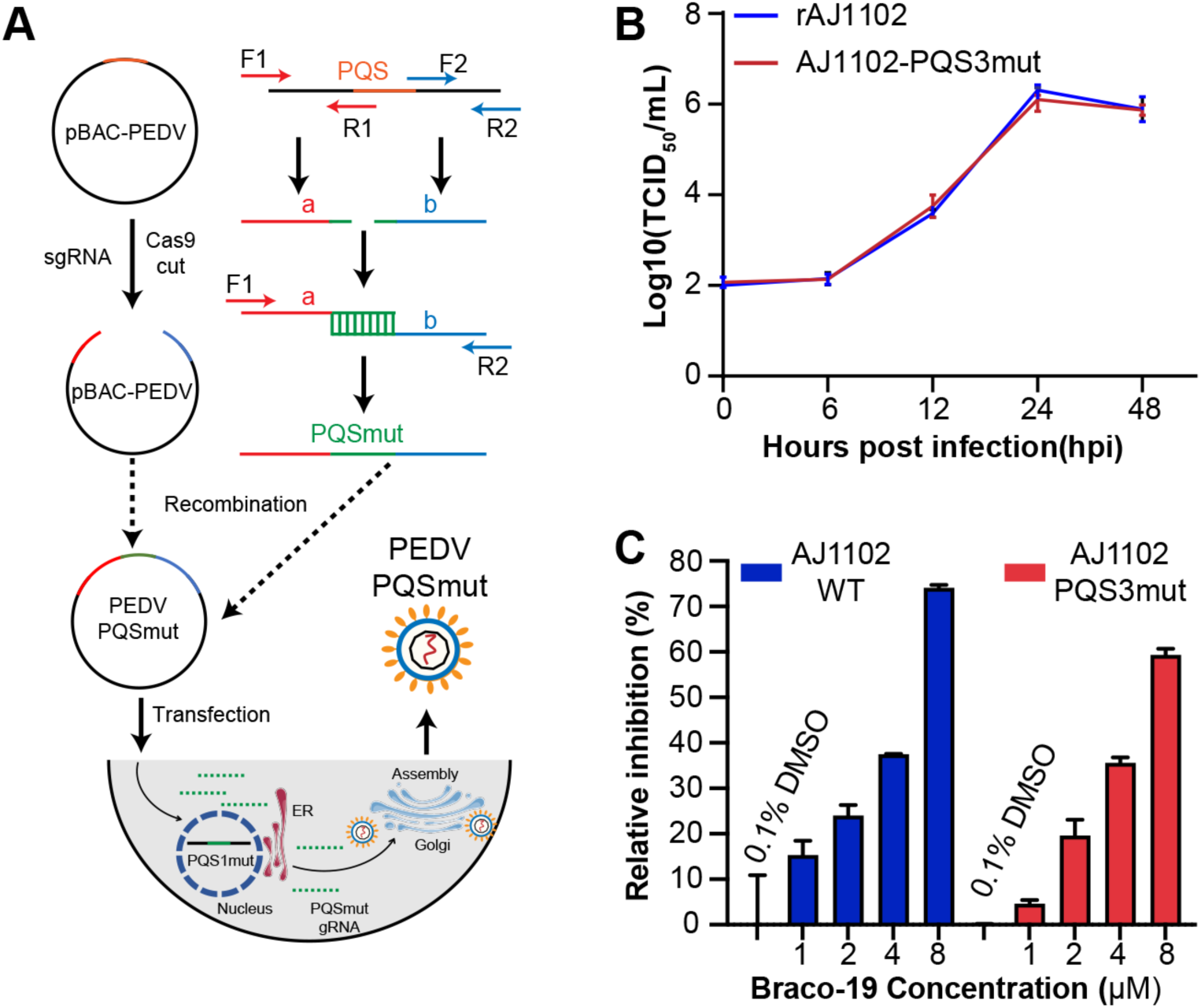
**(A)** Schematic diagram of the process used to obtain the PQSs mutant recombinant strains. **(B)** Proliferation curve of the PQS3 mutant strain. **(C)** The relative inhibition rates of Braco-19 against AJ1102-WT and AJ1102-PQS3mut.

**Figure 4—figure supplement 4.**
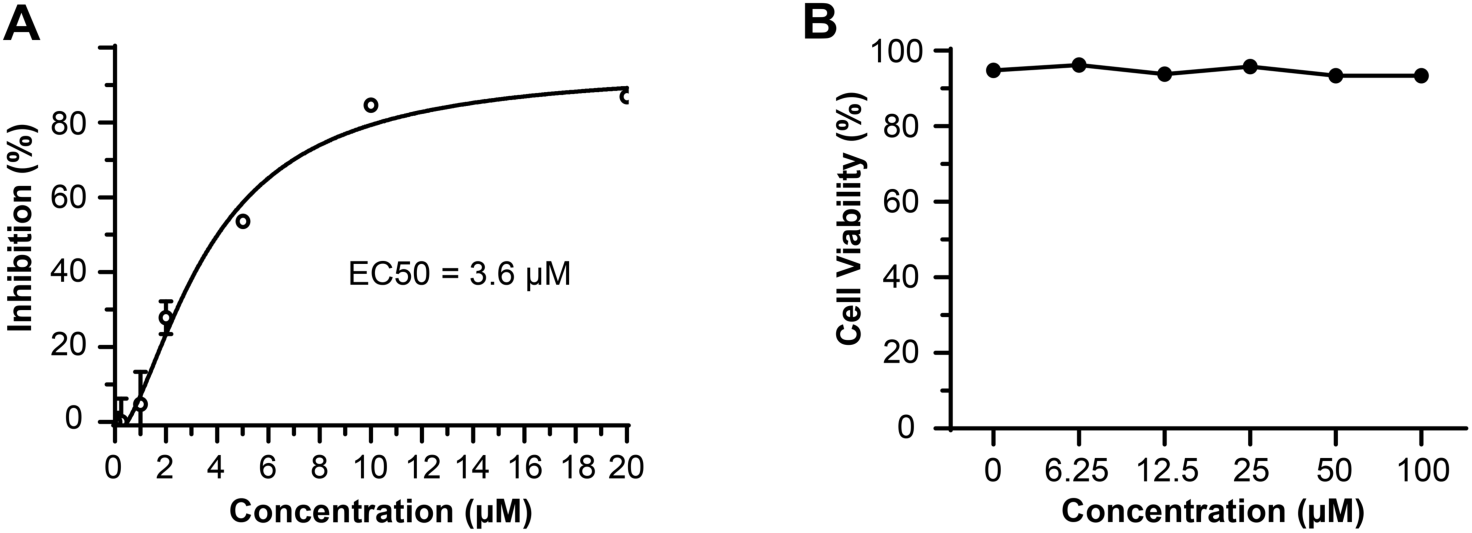
Antiviral activity and cytotoxicity of Braco-19. **(A)** The median effective inhibitory concentration (EC50) of Braco-19 against PEDV AJ1102 in Vero cells and **(B)** the cytotoxicity of Braco-19 for Vero cells.

**Figure 5—figure supplement 1.**
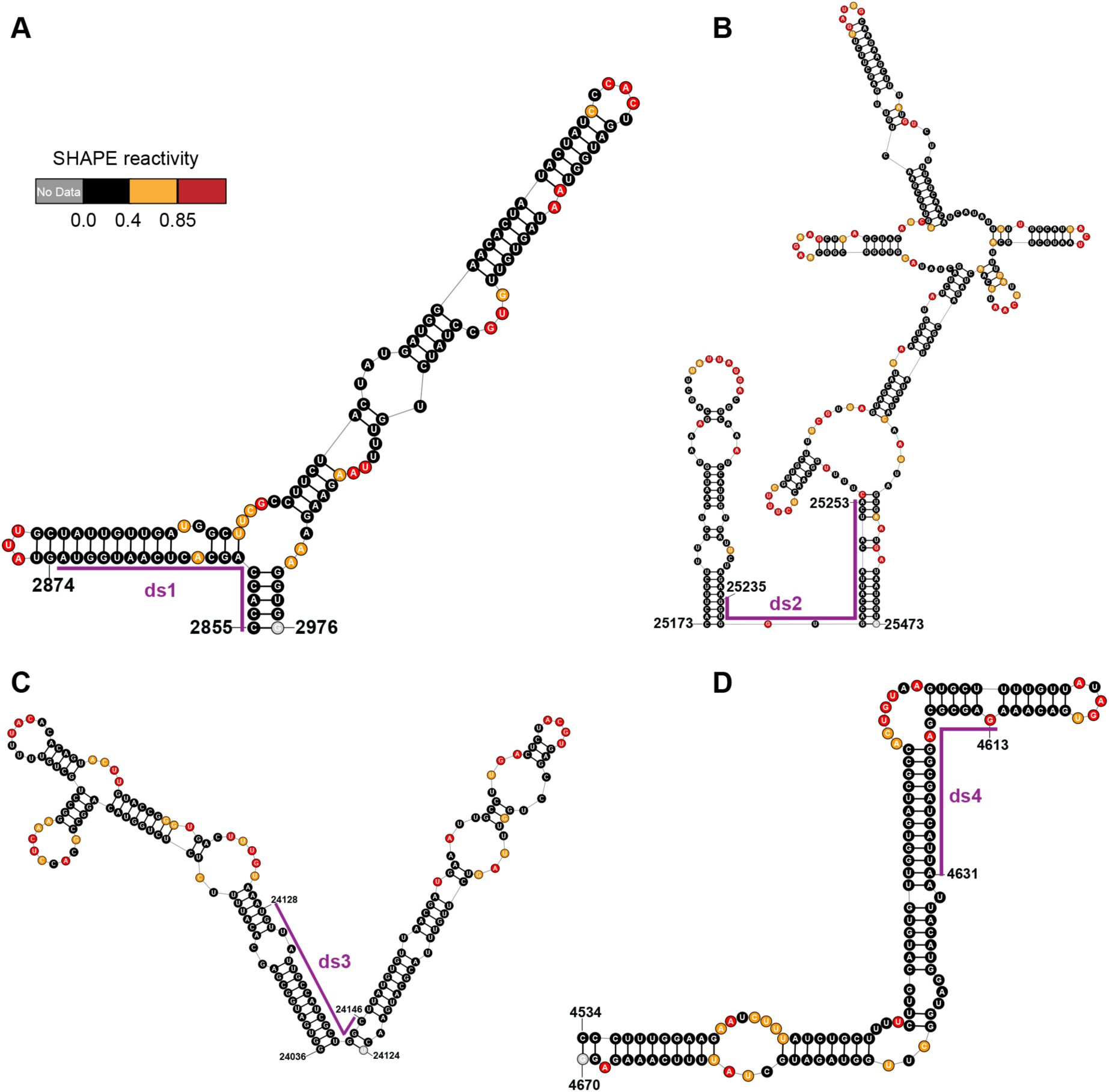
Local secondary structures of the duplex regions targeted by siRNAs. Secondary structure of the **(A)** ds1; **(B)** ds2; **(C)** ds3; **(D)** ds4 target regions predicted by SHAPE reactivity as a constraint. Ds: dual-stranded targeting siRNAs.

**Figure 5—figure supplement 2.**
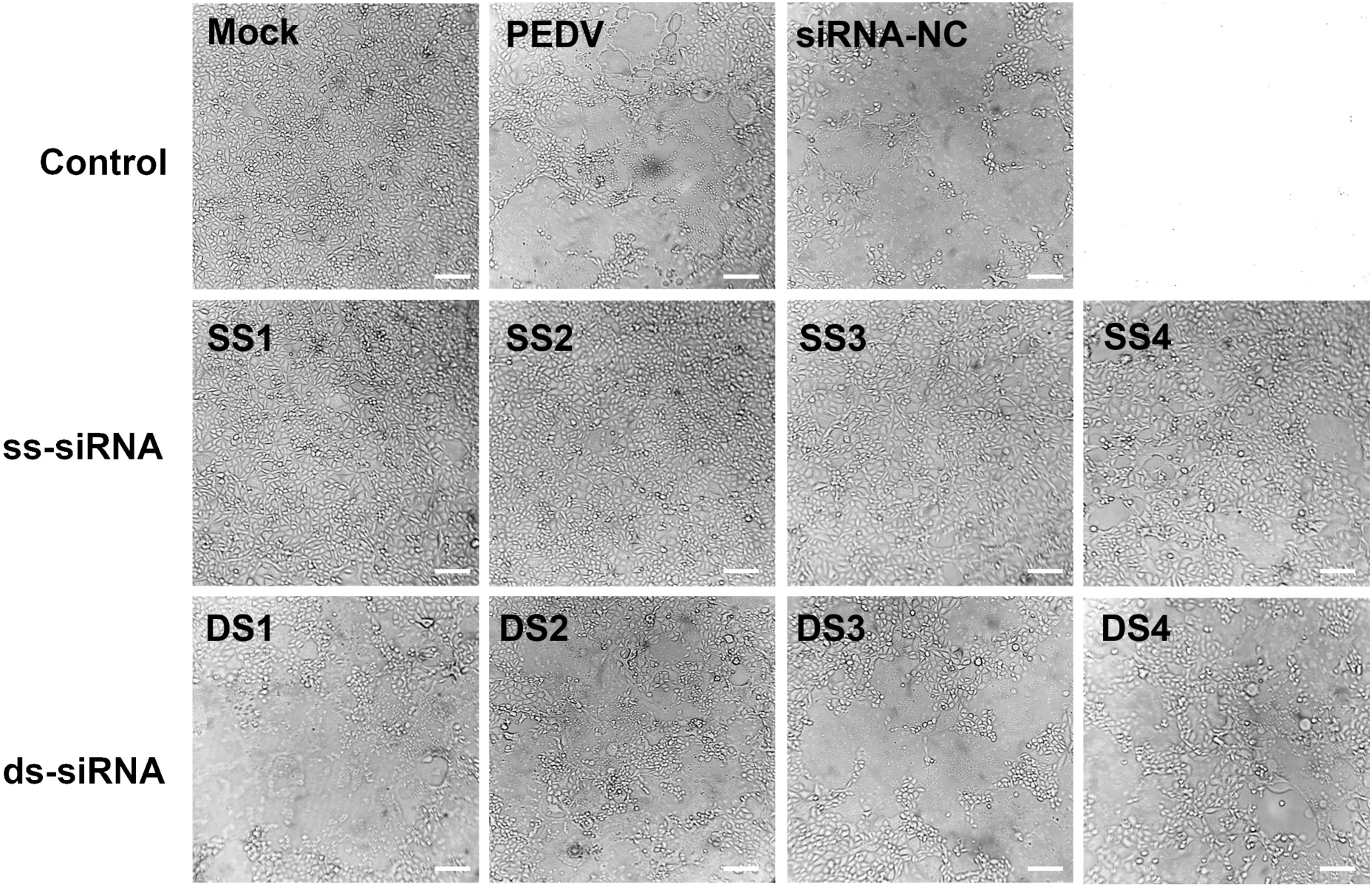
Cytopathic effects (CPE) of siRNA targeting regions of duplex or single-strand on PEDV-infected Vero cells. Scale bar = 50 μm.

