## Supplementary Tables for "Exploiting functional regions in the viral RNA genome as druggable entities"

**Table S1. Oligos used in this study.**

| Oligomer | Sequence (5'-3') |
|---|---|
| PQS1 | GGUUACAGGUGGUUGG |
| PQS1mut | AGUUACAGGUGGUUGG |
| PQS3 | GGCCAAUGGUUCUGGUAAUGG |
| EGFP-PQS1-F | AAGGTTACAGGTGGTTGGGACAGCAAGGGCGAGGAGCT |
| EGFP-PQS1-R | GTCCCAACCACCTGTAACCTTCATGGTGGCGACCGGTAG |
| EGFP-PQS1mut-F | AAAGTTACAGCTGCTTGGGACAGCAAGGGCGAGGAGCTG |
| EGFP-PQS1mut-R | GTCCCAAGCAGCTGTAACTTTCATGGTGGCGACCGGTA |
| PQS1Mut-F1 | CATTTCACAGCACCAAGTTATATGGAGG |
| PQS1Mut-R1 | CCACCTGTAACTTTGATACGATTACCAACA |
| PQS1Mut-F2 | TATCAAAGTTACAGGTGGTTGGGACGATGTTGT |
| PQS1Mut-R2 | ACATCAATGGCCTTTGCTATGCCG |
| PQS3Mut-F1 | ATGGCAAGGTGGTACACGTTA |
| PQS3Mut-R1 | CCATTACCAGAACCATTGGCTAACATCTTA |
| PQS3Mut-F2 | AGCCAATGGTTCTGGTAATGGACAAGCTGT |
| PQS3Mut-R2 | CTTGTTCAAAGGATAACCTGCG |
| Scaffold oligo | AAAAGCACCGACTCGGTGCCACTTTTTCAAGTTGATAACGGACTA GCCTTATTTTAACTTGCTATTTCTAGCTCTAAAAC |
| PQS1m-sgRNA-a | GGATCCTAATACGACTCACTATAGGGACAGCACCAAGTTATATGG GTTTTAGAGCTAGA |
| PQS1m-sgRNA-b | GGATCCTAATACGACTCACTATAGGGATGAGAATTTGTCTCACGG GTTTTAGAGCTAGA |
| PQS3m-sgRNA-a | GGATCCTAATACGACTCACTATAGGGAAGGTGGTACACGTTAAGG GTTTTAGAGCTAGA |
| PQS3m-sgRNA-b | GGATCCTAATACGACTCACTATAGGGCAAACCTTAACAAGAGCGC GTTTTAGAGCTAGA |
| qPCR-nsp12-F | GCTAAGTTGAAGCCAATGCC |
| qPCR-nsp12-R | CCAAATAGTAAAGCCCGCC |
| qPCR-N-F | CCGTGGTGAGCGAATTGAAC |
| qPCR-N-R | GGTTCAGTCTTTGCGCCTTC |
| β-actin-F | CCACCATGTACCCTGGCATT |
| β-actin-R | ACTCCTGCTTGCTGATCCAC |
| Template PQS1 | GGUUACAGGUGGUUGG UCGUA UAGUG AGUCG UAUUA |
| Template PQS1mut | AGUUACAGGUGGUUGG UCGUA UAGUG AGUCG UAUUA |
| FAM-P15 | 5'-FAM-UAAUACGACUCACUA-3' |
| FAM-M17 | 5'-FAM-UAAUACGACUCACUAUA-3' |
| FAM-M20 | 5'-FAM-UAAUACGACUCACUAUACGA-3' |
| FAM-M42 | 5'-FAM- UAAUACGACUCACUAUACGAUUAAUACGACUCACUAUACGAU- 3' |

**Table S2. Characteristics of regions with different SHAPE reactivity and Shannon entropy.**

| Type of the regions | Characteristics | Percentage of the genome | Typical representative regions |
|---|---|---|---|
| Low SHAPE-low Shannon | Base-paired and stable | 26.40% | 5'UTR, M gene |
| High SHAPE-low Shannon | Accessible and stable | 9.59% | TRS-L, FSE |
| High SHAPE-high Shannon | Accessible and dynamic | 6.23% | E gene |
| Low SHAPE-high Shannon | Base-paired and dynamic | 11.65% | 3'UTR |

**Table S3. Location of regions with different SHAPE reactivity and Shannon entropy.**

| Low SHAPE-Low Shannon | | | Low SHAPE-High Shannon | | | High SHAPE-Low Shannon | | | High SHAPE-High Shannon | | |
|---|---|---|---|---|---|---|---|---|---|---|---|
| start | end | length | start | end | length | start | end | length | start | end | length |
| 174 | 227 | 54 | 133 | 172 | 40 | 46 | 75 | 30 | 1595 | 1621 | 27 |
| 372 | 497 | 126 | 253 | 313 | 61 | 318 | 350 | 33 | 2528 | 2562 | 35 |
| 510 | 605 | 96 | 1081 | 1132 | 52 | 791 | 828 | 38 | 2818 | 2844 | 27 |
| 643 | 790 | 148 | 1786 | 1836 | 51 | 1231 | 1256 | 26 | 3090 | 3129 | 40 |
| 983 | 1078 | 96 | 1962 | 2024 | 63 | 1554 | 1573 | 20 | 4396 | 4421 | 26 |
| 1145 | 1190 | 46 | 2348 | 2430 | 83 | 3881 | 3905 | 25 | 5690 | 5734 | 45 |
| 1321 | 1538 | 218 | 2594 | 2650 | 57 | 4463 | 4485 | 23 | 6740 | 6764 | 25 |
| 2111 | 2256 | 146 | 2951 | 3000 | 50 | 4851 | 4873 | 23 | 7195 | 7217 | 23 |
| 2273 | 2347 | 75 | 3135 | 3241 | 107 | 5224 | 5246 | 23 | 8086 | 8112 | 27 |
| 2853 | 2949 | 97 | 3306 | 3439 | 134 | 6617 | 6639 | 23 | 8402 | 8426 | 25 |
| 3700 | 3757 | 58 | 3543 | 3589 | 47 | 9025 | 9045 | 21 | 8557 | 8610 | 54 |
| 4216 | 4299 | 84 | 3804 | 3874 | 71 | 9415 | 9438 | 24 | 8643 | 8684 | 42 |
| 4537 | 4693 | 157 | 3951 | 4038 | 88 | 9817 | 9843 | 27 | 8824 | 8849 | 26 |
| 4699 | 4821 | 123 | 4300 | 4383 | 84 | 10170 | 10227 | 58 | 8880 | 8912 | 33 |
| 4884 | 4938 | 55 | 5319 | 5400 | 82 | 10276 | 10296 | 21 | 9358 | 9404 | 47 |
| 5402 | 5544 | 143 | 6118 | 6361 | 244 | 10571 | 10616 | 46 | 9680 | 9707 | 28 |
| 5585 | 5654 | 70 | 6538 | 6597 | 60 | 10900 | 10923 | 24 | 10516 | 10539 | 24 |
| 5768 | 6117 | 350 | 6669 | 6738 | 70 | 11237 | 11263 | 27 | 10550 | 10570 | 21 |
| 6249 | 6314 | 66 | 6922 | 6967 | 46 | 11291 | 11321 | 31 | 10761 | 10782 | 22 |
| 6442 | 6537 | 96 | 7071 | 7123 | 53 | 11491 | 11520 | 30 | 11098 | 11117 | 20 |
| 6810 | 6859 | 50 | 7237 | 7279 | 43 | 11921 | 11966 | 46 | 12921 | 12946 | 26 |
| 6970 | 7066 | 97 | 7345 | 7410 | 66 | 12060 | 12081 | 22 | 13069 | 13090 | 22 |
| 7412 | 7581 | 170 | 7582 | 7622 | 41 | 12843 | 12862 | 20 | 13721 | 13782 | 62 |
| 7625 | 7678 | 54 | 7813 | 7880 | 68 | 12987 | 13013 | 27 | 14646 | 14673 | 28 |
| 7730 | 7808 | 79 | 9117 | 9234 | 118 | 13037 | 13068 | 32 | 15309 | 15359 | 51 |
| 8710 | 8787 | 78 | 9459 | 9607 | 149 | 13193 | 13247 | 55 | 15482 | 15536 | 55 |
| 9275 | 9337 | 63 | 10310 | 10354 | 45 | 13455 | 13498 | 44 | 16037 | 16083 | 47 |

| | | | | | | | | | | | |
|---|---|---|---|---|---|---|---|---|---|---|---|
| 9761 | 9939 | 179 | 12150 | 12295 | 146 | 13590 | 13617 | 28 | 16996 | 17041 | 46 |
| 10018 | 10101 | 84 | 14008 | 14053 | 46 | 13882 | 13948 | 67 | 18141 | 18197 | 57 |
| 10924 | 11085 | 162 | 15783 | 15829 | 47 | 14058 | 14082 | 25 | 18373 | 18427 | 55 |
| 11663 | 11778 | 116 | 17562 | 17684 | 123 | 14758 | 14806 | 49 | 18631 | 18685 | 55 |
| 12095 | 12149 | 55 | 17925 | 17964 | 40 | 15676 | 15706 | 31 | 18751 | 18771 | 21 |
| 12312 | 12589 | 278 | 19218 | 19291 | 74 | 16160 | 16181 | 22 | 19309 | 19346 | 38 |
| 12659 | 12828 | 170 | 19894 | 19940 | 47 | 16333 | 16360 | 28 | 20141 | 20185 | 45 |
| 13262 | 13319 | 58 | 21201 | 21361 | 161 | 16595 | 16630 | 36 | 20705 | 20741 | 37 |
| 13513 | 13564 | 52 | 22431 | 22477 | 47 | 16828 | 16920 | 93 | 20805 | 20828 | 24 |
| 14327 | 14493 | 167 | 24312 | 24420 | 109 | 17043 | 17149 | 107 | 20873 | 20902 | 30 |
| 14700 | 14751 | 52 | 24737 | 24775 | 39 | 17162 | 17185 | 24 | 21094 | 21123 | 30 |
| 14862 | 14903 | 42 | 24835 | 24884 | 50 | 17288 | 17326 | 39 | 21266 | 21324 | 59 |
| 14932 | 15004 | 73 | 26838 | 26924 | 87 | 17432 | 17466 | 35 | 21375 | 21403 | 29 |
| 15719 | 15782 | 64 | 27723 | 27790 | 68 | 18215 | 18235 | 21 | 22351 | 22418 | 68 |
| 15861 | 15909 | 49 | 27874 | 27935 | 62 | 18707 | 18750 | 44 | 22527 | 22558 | 32 |
| 16179 | 16287 | 109 | | | | 19016 | 19060 | 45 | 23639 | 23688 | 50 |
| 16521 | 16702 | 182 | | | | 19128 | 19163 | 36 | 23936 | 23957 | 22 |
| 17774 | 17924 | 151 | | | | 19482 | 19507 | 26 | 24427 | 24464 | 38 |
| 18043 | 18111 | 69 | | | | 19560 | 19619 | 60 | 25491 | 25521 | 31 |
| 18233 | 18282 | 50 | | | | 19645 | 19700 | 56 | 25612 | 25632 | 21 |
| 18825 | 18877 | 53 | | | | 20501 | 20533 | 33 | 26419 | 26447 | 29 |
| 19076 | 19126 | 51 | | | | 21404 | 21425 | 22 | 27493 | 27515 | 23 |
| 19402 | 19455 | 54 | | | | 21468 | 21571 | 104 | | | |
| 19717 | 19768 | 52 | | | | 21739 | 21800 | 62 | | | |
| 19875 | 20115 | 241 | | | | 21970 | 22008 | 39 | | | |
| 20321 | 20442 | 122 | | | | 22088 | 22148 | 61 | | | |
| 20944 | 20985 | 42 | | | | 22230 | 22277 | 48 | | | |
| 21689 | 21731 | 43 | | | | 22708 | 22753 | 46 | | | |
| 21851 | 21922 | 72 | | | | 22773 | 22805 | 33 | | | |
| 22931 | 22989 | 59 | | | | 22902 | 22927 | 26 | | | |
| 23113 | 23175 | 63 | | | | 23044 | 23086 | 43 | | | |
| 23404 | 23457 | 54 | | | | 23198 | 23257 | 60 | | | |
| 23490 | 23559 | 70 | | | | 23321 | 23381 | 61 | | | |
| 24016 | 24091 | 76 | | | | 23471 | 23494 | 24 | | | |
| 24103 | 24280 | 178 | | | | 23754 | 23796 | 43 | | | |
| 24777 | 24960 | 184 | | | | 23993 | 24012 | 20 | | | |
| 25186 | 25266 | 81 | | | | 24619 | 24642 | 24 | | | |
| 25291 | 25396 | 106 | | | | 25134 | 25167 | 34 | | | |
| 25713 | 25846 | 134 | | | | 25457 | 25477 | 21 | | | |
| 25902 | 25985 | 84 | | | | 25510 | 25534 | 25 | | | |
| 26036 | 26115 | 80 | | | | 25672 | 25692 | 21 | | | |
| 26145 | 26263 | 119 | | | | 26449 | 26492 | 44 | | | |
| 26328 | 26399 | 72 | | | | 26522 | 26547 | 26 | | | |
| 26666 | 26731 | 66 | | | | 26945 | 26967 | 23 | | | |
| 27093 | 27235 | 143 | | | | 27517 | 27542 | 26 | | | |
| 27268 | 27398 | 131 | | | | 27677 | 27712 | 36 | | | |
| 27553 | 27665 | 113 | | | | | | | | | |

**Table S4. Binding affinities of Compounds with 5'UTR-SL5 in the PEDV genome.**

| Compounds | Names | structural formula | Binding affinities |
|---|---|---|---|
| 1 | 1-(2-methylpyrimidin-4-yl)piperidin-4-amine | 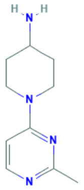 | 4.75 µM |
| 2 | 3,4-Dimethoxybenzylamine | 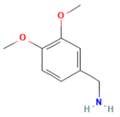 | 0.11 mM |
| 3 | N-ethyl-1-(pyridin-4-yl)piperidin-4-amine | 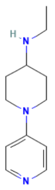 | 18.8 µM |
| 4 | 4-(4-Chlorophenyl)pyrimidin-2-amine | 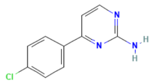 | 1.86 µM |
| 5 | 5-Methoxybenzo[d]thiazol-2-amine | 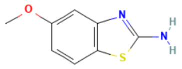 | -- |
| 6 | N-3-pyridinylpyrazolo[1,5-a]pyrimidine-3-carboxamide | 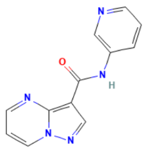 | 1.10 mM |
| 7 | N-methyl-1-(1-phenylpyrazol-4-yl)methanamine | 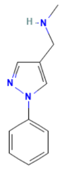 | 13.3 µM |

**Table S5. PQSs with high conservation in the PEDV genome.**

| number | PQSs | position | length | conservation | location | Characteristics of the region |
|---|---|---|---|---|---|---|
| PQS1 | GGUUACAG GUGGUUGG | 3109 | 16 | 98.37% | nsp3 | high SHAPE-high Shannon |
| PQS2 | GGUAUUGG UGGUGAGC GG | 10988 | 18 | 97.63% | nsp6 | low SHAPE-low Shannon |
| PQS3 | GGCCAAUG GUUCUGGU AAUGG | 12322 | 21 | 98.52% | nsp10 | low SHAPE-low Shannon |
| PQS4 | GGUUGUUG GCUGGCUA AUGG | 12545 | 20 | 93.79% | nsp10 | low SHAPE-low Shannon |
| PQS5 | GGGCUGGU GGUUUGG | 17720 | 15 | 98.82% | nsp14 | Others |
| PQS6 | GGUGGUAU | 23508 | 17 | 92.16% | S gene | low SHAPE-low |

| | | | | | | |
|---|---|---|---|---|---|---|
| | GGUGCUAG G | | | | | Shannon |
| PQS7 | GGCCGUGG UGGGUUUG G | 24607 | 17 | 98.22% | S gene | Others |

**Table S6: Genomic locations and structural features of PQS-long chain regions**

| PQS-long chain | Start | End | Length | Structural characteristics |
|---|---|---|---|---|
| PQS1 | 3091 | 3276 | 186 nt | high SHAPE-high Shannon |
| PQS2 | 10924 | 11087 | 164 nt | Low SHAPE-low Shannon |
| PQS3 | 12200 | 12590 | 391 nt | Low SHAPE-low Shannon |
| PQS4 | 12336 | 12588 | 253 nt | Low SHAPE-low Shannon |
| PQS6 | 23429 | 23536 | 108 nt | Low SHAPE-low Shannon |

**Table S7. Sequence and characterisation of target regions of siRNAs.**

| Target region | siRNA | Target Sequences | Local | Conservation | Average SHAPE reactivity | Number of unpaired bases |
|---|---|---|---|---|---|---|
| Single-Strand | ss-1 | UCAAUUCAACU AAACGAAA | 5'UTR (58-76 nt) | 88.96% | 0.90 | 15 |
| | ss-2 | GGCUUAUAAGU CCGUUUUU | Nsp3 (4462-4480 nt) | 97.37% | 0.54 | 10 |
| | ss-3 | GUACCCUAUUA UUGUUUUC | S (21738-21756 nt) | 86.90% | 0.81 | 13 |
| | ss-4 | GGCUGUUUUUA UUUCUCCU | Nsp13 (16899-16917 nt) | 92.25% | 0.44 | 12 |
| Dual-Strand | ds-1 | CCACCAGCACU CAAUGGUA | Nsp2 (2855-2873 nt) | 92.38% | 0.08 | 2 |
| | ds-2 | GGUGGUGACCA UUACAUCA | ORF3 (25235-25253 nt) | 90.01% | 0.16 | 2 |
| | ds-3 | UGUUAUUGCCA UCGCUGGC | S (24128-24146 nt) | 88.96% | 0.09 | 3 |
| | ds-4 | GAGCGCGAGGC GAUCAUUA | Nsp3 (4613-4631 nt) | 95.40% | 0.27 | 3 |
| Non-target | siRNA- | UCCAGAUCUGU | -- | -- | -- | -- |

| | | |
|---|---|---|
| control | NC | AAGGGUAC |

**Table S8: Structural features of anti-SARS-CoV-2 siRNA target regions.**

| siRNA(Bowden-Reid et al., 2023) | Sequence (Sense) | target region | high SHAPE-low Shannon(Manfredonia et al., 2020) |
|---|---|---|---|
| 2 | 5`-CUUCCCAGGUAACAAACCAdTdT-3` | 5`UTR (16-34 nt) | Yes |
| 7 | 5`-CGUCCGGGUGUGACCGAAAdTdT-3` | 5`UTR (241-259 nt) | No |
| 16 | 5`-GUAGUACUUUCUUUUGAACdTdT-3` | Spike (23093-23111 nt) | No |
| 18 | 5`-GCUACAUCACGAACGCUUUdTdT-3` | Membrane (27033-27051 nt) | Yes |
| 21 | 5`-GCCAUCCUUACUGCGCUUCdTdT-3` | Envelope (26338-26356 nt) | No |
| 25 | 5`-GGGUUGCAACUGAGGGAGCdTdT-3` | Nucleocapsid (28668-28696 nt) | Yes |
| 27 | 5`-CGAGAAAACACACGUCCAAdTdT-3` | ORF1ab (NSP1, 292-310 nt) | Yes |
| 30 | 5`-GGCAUUCAGUACGGUCGUAdTdT-3` | ORF1ab (NSP1, 545-563 nt) | Yes |
